# RBM25 exhibits distinct RNA- and chromatin-associated target programs in human embryonic stem cells

**DOI:** 10.64898/2026.10.05.756609

**Authors:** Yuanpu Sun, Tamar Lahav, Fabian Glaser, Tamar Hashimshony, Yael Mandel-Gutfreund

## Abstract

RNA-binding proteins (RBPs) are increasingly recognized to perform chromatin-associated functions in addition to their canonical roles in RNA regulation, yet how these activities are coordinated in human embryonic stem cells (hESCs) remain poorly understood. Here, we characterize distinct RNA- and chromatin-associated activities for RBM25 in hESCs. Transcriptome-wide profiling showed that RBM25 preferentially associates with transcripts involved in RNA processing and splicing, consistent with its canonical role as an RBP involved in post-transcriptional regulation. Genome-wide chromatin profiling further uncovered a distinct chromatin-associated program, with RBM25 occupying accessible promoters of actively expressed cell-cycle genes and introns predominantly associated with transcriptionally inactive neuronal genes. Interestingly, RBM25 chromatin-binding sites showed substantial co-occupancy with regions bound by the transcription factor CEBPZ in hESCs. We further show that RBM25 and CEBPZ physically interact and exhibit shared chromatin-binding patterns at both promoter and intronic regions. Importantly, reciprocal depletion experiments revealed that RBM25 is required for CEBPZ occupancy at CCAAT-containing promoters of highly expressed cell-cycle genes, whereas CEBPZ is not similarly required for RBM25 chromatin occupancy. Together, our findings establish RBM25 as a multifunctional factor with distinct roles in RNA processing and chromatin-associated regulation and reveal a context-dependent RBP–TF interaction that contributes to transcriptional regulation in hESCs.

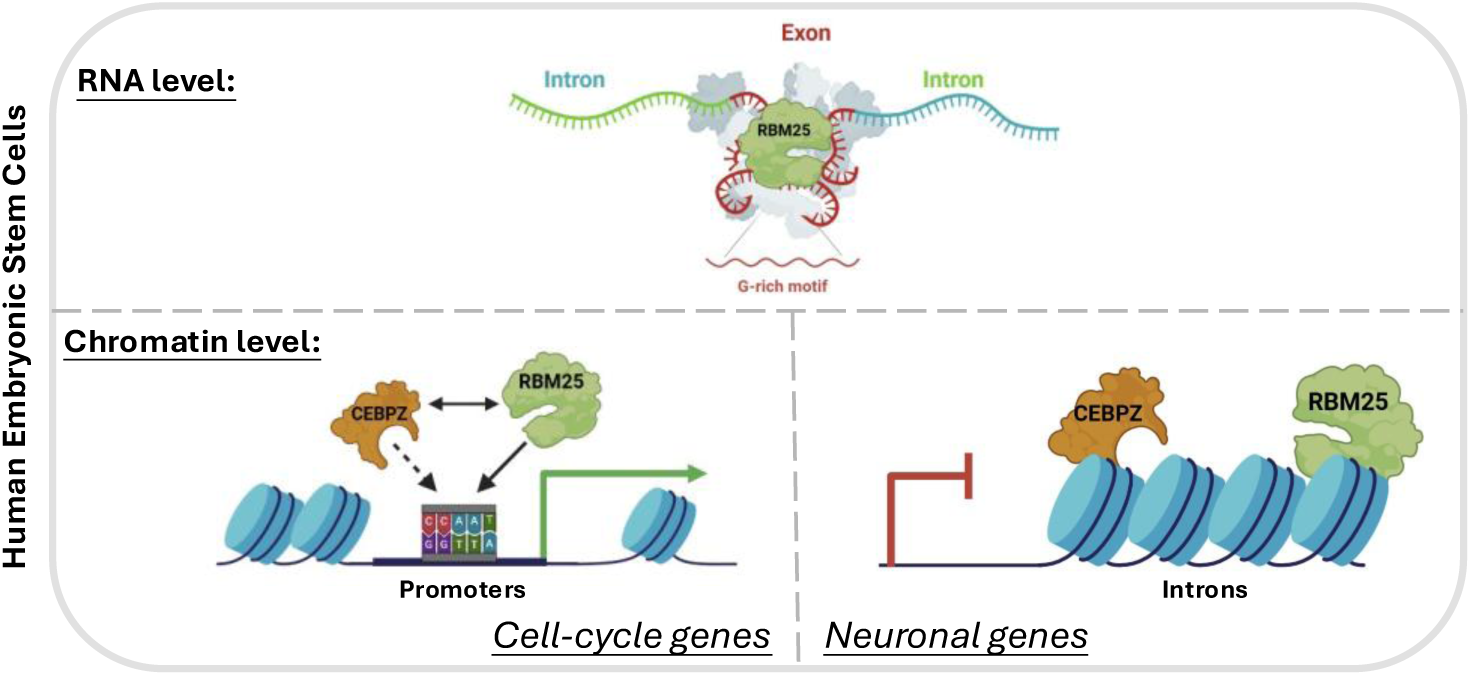

## Introduction

The unique ability of embryonic stem cells (ESCs) to self-renew and differentiate to all three germ layers is maintained through tightly coordinated regulatory mechanisms at multiple levels of gene expression (1–5). Among these, RNA-binding proteins (RBPs) have also been shown as key regulators of pluripotency and cell state transitions in ESCs (6). RBPs function primarily through interactions with RNA to form ribonucleoprotein complexes (RNPs), thereby regulating diverse post-transcriptional processes, including pre-mRNA splicing, RNA modification, stability, and translation. Some RBPs have been shown to play critical roles in maintaining pluripotency (6, 7). For example, LIN28 has a well-established role in pluripotency via post-transcriptional regulation of the pluripotency-associated factor OCT4, and in suppressing let-7 microRNA activity to maintain the pluripotent state in ESCs (8–10). Also, several splicing factors such as FOX2 and SON have been shown to regulate pluripotency-specific alternative splicing in human ESCs (hESCs) (11, 12). In recent years, advances in high throughput RNA-binding technologies, such as UV crosslinking-based RNA interactome capture (RIC) have enabled the systematic identification of RBPs, uncovering numerous previously uncharacterized RBPs in ESCs. Many of these RBPs are preferentially expressed in ESCs relative to differentiated cells and have been implicated in the regulation of pluripotency and differentiation (13, 14). Together, these findings highlight the central role of RBPs in governing pluripotency and early differentiation in ESCs (7).

The RNA interactome studies in ESCs have also identified RBPs annotated with additional functions including roles in transcription related functions (14, 15). Accumulating evidence indicates that a subset of RBPs can associate with chromatin and actively participate in transcriptional control, in addition to their canonical RNA-binding activities (16–20). For example, RBPs are shown to interact with nascent RNA or RNA polymerase II (Pol II) to affect the pause, elongation, and termination of RNA Pol II during transcription (21–25). Furthermore, some RBPs directly bind chromatin and recruit transcriptional regulators to modulate gene expression (19, 26, 27). However, the chromatin binding of RBPs can also contribute to its post-transcription regulation functions. For example, U2AF2 is shown to bind H3K36me3 marked chromatin and regulate co-transcriptional splicing (28). Conversely, certain DNA binding proteins (DBPs) have been shown to interact with RNA and regulate RNA processing and gene expression (29–32). Those proteins with capacity to bind both DNA and RNA, are defined as DNA- and RNA- binding proteins (DRBPs) (33), can likely contribute to the cross-talk between transcriptional and post-transcriptional regulation. In practice, DRBPs are typically identified by integrating genome-wide DNA-binding assays, such as ChIP-seq or CUT&RUN, with RNA-binding approaches, including CLIP-seq or RNA interactome capture coupled with mass spectrometry (20, 34, 35). Based on their functional properties, DRBPs can be broadly classified into three categories: 1, DRBPs that competitively bind to DNA and RNA. 2, DRBPs that have dual function on their DNA and RNA targets. 3, DRBPs that bind to DNA and RNA simultaneously (33). This emerging group of DRBPs highlights an integrated framework of gene regulation that couples transcriptional and post-transcriptional processes.

Moreover, RBPs can also regulate gene expression through interactions with transcription factors (TFs). These interactions are often enriched at active promoters (19) and can occur through multiple mechanisms, including TF-driven recruitment of RBPs, RBP-mediated recruitment of TFs, or cooperative binding of both factors (19, 36–38). Functional interplay between TFs and RBPs is frequently reflected by shared DNA-binding motifs, overlapping chromatin occupancy, and physical interactions detectable by biochemical assays such as co-immunoprecipitation (co-IP). The published examples of functional interactions between RBPs and TFs demonstrate diverse strategies for coupling transcriptional activity with downstream RNA processing, such interactions provide an additional mechanism for coupling transcriptional activity with downstream RNA processing, thereby coordinating multiple layers of gene regulation. Despite these advances, the mechanisms by which RBPs are recruited to specific genomic loci and functionally cooperate with TFs in hESCs remain largely unknown.

RBM25 is a well-characterized RBP with established roles in pre-mRNA splicing across multiple cell types, including acute myeloid leukemia (AML) and colon cancer cells (39–43). It contains multiple functional regions and domains, including an RNA recognition motif (RRM), a glutamate/aspartate-rich (RE/RD-rich) region involved in protein–protein interactions (39), and a PWI domain capable of binding both single- and double-stranded nucleic acids (44). While most studies have focused on its role in RNA processing, emerging evidence suggests that RBM25 may also participate in chromatin-associated regulation. Specifically, RBM25 has been shown to interact with the TF YY1 to facilitate promoter–enhancer interactions in HepG2 cells (19). In addition, RBM25 has been identified in nuclear RNA interactomes in hESCs (14) and is highly expressed in mouse ESCs, where it contributes to the regulation of pluripotency-associated gene expression and epigenetic states (45). These observations raise the possibility that RBM25 has a key role in linking transcriptional and post-transcriptional regulation in human and mouse. However, its specific function in hESCs, particularly as a DRBP in the context of chromatin association and TF interaction has not been studied.

In this study, we investigate the function of RBM25 in hESCs by integrating genome-wide chromatin profiling, transcriptomic and proteomic analyses. We demonstrate that RBM25 associates with both RNA and chromatin via different target genes, supporting its function as a dual functional DNA- and RNA-binding protein in hESCs. At the chromatin level, RBM25 exhibits distinct binding patterns: at promoter regions, it preferentially associates with CCAAT motifs in actively expressed cell cycle genes, whereas within intronic regions, it binds within closed chromatin regions associated with silenced neuronal genes. Furthermore, we identify the TF CEBPZ as the potential interacting partner of RBM25. We show that RBM25 and CEBPZ interact physically and share co-binding genes at both promoter and intronic regions. Notably, RBM25 is required for efficient CEBPZ binding at promoters of cell cycle genes, suggesting a coordinated regulatory mechanism. Together, our results establish RBM25 as a multifunctional RNA- and chromatin-associated regulator in hESCs and reveal a promoter-specific role in which RBM25 supports CEBPZ occupancy at CCAAT-containing cell-cycle gene promoters.

## Materials and methods

### co-occupancy analysis of TF-RBP pairs

TF ChIP–seq datasets were obtained from the ENCODE (GRCh38) database. Chip-seq datasets were included in the analysis only if they (i) were generated under native conditions without epitope tagging, chemical treatment, or genetic modification, (ii) used validated antibodies, and (iii) profiled proteins previously classified as DNA-binding factors or sequence-specific transcription factors. These criteria yielded 136 TF datasets. RBP ChIP–seq datasets were obtained from a previous ENCODE-based study that curated high-quality RBP ChIP–seq datasets, resulting in 20 RBPs chip-seq datasets profiled in K562 cells (19).

Promoter regions were defined as ±1 kb around annotated transcription start sites (TSSs) using the hg38 annotation. Only promoter-associated peaks were retained for subsequent analyses. Proteins with fewer than 800 promoter-associated peaks were excluded because of insufficient statistical power. For proteins with more than 1, 000 promoter-associated peaks, the top 1, 000 peaks ranked by peak score were retained to minimize bias and to standardize peak numbers across datasets. After filtering, 127 TFs and 15 RBPs were included in the co-occupancy analysis.

Peak overlap was quantified using the GenomicRanges package in R (46). For each TF–RBP pair, the observed overlap count was defined as the number of TF-RBP promoter peaks overlapping at least 1bp. To assess whether the observed overlap exceeded random expectation, a permutation-based background distribution was generated. The promoter-associated peaks from all unfiltered 136 TF datasets were pooled to create a background peak universe. From this pooled set, 1, 000 peaks were randomly sampled to generate a background containing 1000 peak sets. For each RBP, the number of overlapped background peaks was calculated for each permutation, generating an RBP-specific null distribution.

To evaluate the significance of TF–RBP co-occupancy, the observed overlap count for each TF–RBP pair was compared with the corresponding RBP-specific background distribution. The z-score was calculated based on the difference between the observed overlap count and the mean overlap of the 1, 000 randomly generated background peak sets, normalized by the standard deviation of the background distribution.

### Cell culture

hESCs (TE03) were maintained in suspension culture in YF100 medium (82% DMEM/F12, 15% KnockOut Serum replacement, 0.1mM 2-mercaptoethanol, 100 ng/ml FGF basic (b/FGF), 1% NEAA, 3 mM L-glutamine, and Pen/Strep). For adherent culture, hESCs were dissociated into single cells using EGTA followed by TrypLE treatment and seeded onto Matrigel-coated plates in mTeSR1 medium (stemcell technologies) supplemented with 5μM ROCK inhibitor (RI). Cells were subsequently maintained in mTeSR1 medium containing RI.

### seCLIP-seq

RBM25 seCLIP was performed in TE03 hESCs as previously described (47), with a few alterations. The duplicates were performed in parallel; cells were crosslinked under 254-nm UV light at 3.2 J/cm^2^. Approximately 30 million cells per sample were lysed in the presence of protease inhibitors (PI) and sonicated using a Covaris M220 with settings of I: 75, B: 200, and D:5 for 2min. Then, the genomic DNA and RNA were partially digested by Turbo DNase (Invitrogen) for 5 min and RNase I for 3 min at 37℃. The protein-RNA complexes were isolated by centrifugation. Pre-clearing was performed with lysis buffer washed protein-G beads (Invitrogen 10003D) at 4℃ for 30min. The immunoprecipitation was performed by incubating with the anti-RBM25 antibody (Abcam ab72237) for 4 hours followed by 2 hours incubation with protein-G beads. After washing, end repair, and 3’ adaptor ligation, the protein-RNA complexes were eluted and resolved by denaturing gel electrophoresis. The complexes were then transferred onto a nitrocellulose membrane. After cutting and isolating the RBM25 bound area, the membrane-bound RNA was purified with proteinase K incubation for 1 h at 50°C. The released RNA was then purified by acidic phenol/chloroform extraction and concentrated using RNA Clean & Concentrator-5 kit (Zymo research). The cDNAs were synthesized using SuperScript III reverse transcriptase (Invitrogen). After cleaning and adapter ligation, the libraries were amplified by PCR and size selection was performed on agarose gel. The libraries were subjected to single-end sequencing on NextSeq 2000 platform.

### seCLIP analysis

The sequencing raw data of RBM25 seCLIP was analyzed according to published pipeline(48). The raw data processing was done for all IP and SMI samples. Adapter sequences were trimmed using Cutadapt v3.5 (49), and unique molecular identifiers (UMIs) were extracted using UMI-tools (50). Reads were aligned to the human reference genome (GRCh38) using STAR v2.7.11b (51) and sorted, filtered by SAMtools V1.13 (52). The PCR duplicates were removed using UMI-tools.

The two mapped and UMI-deduplicated IP .bam files were merged and sorted and used for the cross-linking peak calling with the PureCLIP tool (48) on the Galaxy Europe server (https://usegalaxy.eu). The identified cross-linking regions were used for creating coverage counts for each IP and SMI samples using BEDTools (53) and normalized by counts per million. The differential analysis between IP and SMI samples was done by NOISeq package(54) in R. The significant peaks were defined as probability (P) > 0.85 and log2Fold-Change (M) > 1.5. Peaks were annotated using GENCODE release v48 (55). Peaks overlapping pseudogenes, tRNAs, rRNAs, or transcripts annotated as TEC (To be Experimentally Confirmed) were excluded from downstream analyses. For de novo motif discovery, significant peaks were extended by 30 bp on each side and analyzed using XSTREME from the MEME Suite (https://meme-suite.org/meme/). Gene region binding distributions and exon metagene profiles were generated using GenomicPlot (https://github.com/shuye2009/GenomicPlot). Gene Ontology enrichment analysis was performed using ShinyGO (https://bioinformatics.sdstate.edu/go/). For visualization in the UCSC Genome Browser (56), deduplicated IP and SMI BAM files were converted to CPM-normalized BigWig files using the bamCoverage function in deepTools (57). RBM25-binding sites containing the identified de novo motif were detected using RBPmap (58).

### CUT&RUN following siRNA knockdown

CUT&RUN following siRNA-mediated knockdown was performed as previously described (59–61) with minor modifications. Approximately 100, 000 cells transfected with target-specific or negative control siRNAs were used for each reaction. Cell pellets were resuspended in nuclei extraction (NE) buffer (20 mM HEPES-KOH, pH 7.9, 10 mM KCl, 0.5 mM spermidine, 0.1% Triton X-100, 20% glycerol) supplemented with PI and incubated on ice for 10 min to isolate nuclei. The cytoplasmic fraction was used for RNA extraction using AMPure XP beads. Nuclei were immobilized on Concanavalin A-coated magnetic beads in binding buffer (20mM HEPES-KOH pH7.9, 10mM KCl, 1mM CaCl2, 1mM MnCl2) for 10min at 4 °C with rotation. Bead-bound nuclei were washed twice with wash buffer (20mM HEPES-NaOH pH 7.5, 150mM NaCl, 0.5mM spermidine, 0.1% BSA) containing EDTA and incubated with anti-RBM25 (Abcam ab72237), anti-CEBPZ (proteintech 25612-1-AP), or normal rabbit IgG ‘negative control’ for 2 h at 4 °C with rotation. Following antibody incubation, nuclei were washed and incubated with pAG/MNase fusion protein (Addgene plasmid #123461) for 1 h at 4 °C. Chromatin digestion was initiated by the addition of CaCl₂ and allowed to proceed for 30 min at 0.1 °C. Reactions were terminated by adding STOP buffer (100 mM EGTA and 125 μg/mL RNase A). DNA fragments were released by proteinase K digestion and purified using SILANE bead cleanup. Sequencing libraries were prepared using the NEBNext Ultra II DNA Library Prep Kit, purified with AMPure XP beads, and paired-end sequencing was performed on an Illumina NextSeq 2000 platform.

### CUT&RUN analysis

CUT&RUN sequencing data were processed using a standard analysis pipeline (59). Adapter sequences were trimmed using Trimmomatic v0.39 (62), and reads were aligned to the human reference genome (GRCh38) using Bowtie2 (63). Reads mapping to ENCODE blacklist regions were removed using BEDTools, and BAM files were sorted and indexed using SAMtools. Genome-wide coverage tracks (BedGraph) were generated using BEDTools. RBM25 and CEBPZ peaks were called using SEACR (64), with the corresponding IgG CUT&RUN bedGraph tracks used as background controls. Enriched motifs were detected and visualized using Homer (65). The differential binding analysis before and after knock down was done by DESeq2 R pachage (66).

### Cell nuclei isolation

Cell nuclei were isolated as previously described (67) with minor modifications. Approximately 10 million hESCs were washed three times with cold PBS. Subsequently, the cells were resuspended in 4.8ml hypotonic lysis buffer (10mM HEPES pH 7.6, 10mM KCl, 0.1mM EGTA, and 0.1mM EDTA) with 0.5mM DTT and PI and rotated for 10min at 4°C. 144ul IGEPAL was added and mixed with the cell suspension and incubated on ice for 10min. Then the cells were homogenized by passing through syringe with 21-gauge needle. The isolated nuclei were pelleted by centrifugation (14, 000×g, 30s, 4°C), washed three times with hypotonic buffer containing 0.5mM DTT, and used immediately or stored at −80°C.

### Co-Immunoprecipitation

Approximately 10 million hESCs or isolated nuclei were lysed in 1 mL co-IP lysis buffer (20 mM Tris HCl, pH 8.0, 2 mM EDTA, 150 mM NaCl, 1% Triton X-100, 0.1% SDS), supplemented with PI and DNase. Lysates were incubated for 30 min at 4 °C with rotation and clarified by centrifugation at 12, 000 rpm for 20 min at 4 °C. The supernatants were pre-cleared by incubation with normal IgG for 1 h, followed by protein G beads for an additional 1 h at 4 °C with rotation. Pre-cleared lysates were incubated with anti-RBM25 (Abcam ab72237) or anti-CEBPZ (Bethyl A303-153A) antibodies for 6 h at 4 °C, followed by incubation with protein G beads for 1 h. Beads were washed three times with lysis buffer.

For western blot analysis, bound proteins were eluted with 1× NuPAGE loading buffer at 50 °C for 10 min with shaking. For mass spectrometry analysis, beads were washed an additional three times with ice-cold PBS prior to downstream sample preparation.

### Western Blot

Protein samples or immunoprecipitated complexes were mixed with 1× NuPAGE loading buffer supplemented with 10 mM DTT and denatured at 95 °C for 5 min. Samples were separated on 8% SDS–PAGE gels and transferred to PVDF membranes. Membranes were probed with anti-RBM25 (Abcam ab72237, 1:2, 500), anti-CEBPZ (Bethyl A303-153A, Proteintech 25612-1-AP, 1:5, 000), anti-Tubulin (Abcam ab7291, 1:5000) and anti-GAPDH (Abcam ab8245, 1:10000) antibodies according to standard western blotting procedures.

### Proteomics analysis

The proteins on the beads were reduced in 3mM DTT, 100mM ammonium bicarbonate and 8M Urea, modified with 10mM iodoacetamide in 100mM ammonium bicarbonate and 8M Urea (room temperature 30 min in the dark) and digested in 2M Urea, 25mM ammonium bicarbonate with modified trypsin (Promega), overnight at 37°C in a 1:50 (M/M) enzyme-to-substrate ratio. The tryptic peptides were desalted using Oasis HLB 96-well µElution Plate (Waters), dried and re-suspended in 0.1% Formic acid in 2% acetonitrile.

The resulting peptides were analyzed by LC-MS/MS using a Q Exactive HFX mass spectrometer (Thermo) fitted with a capillary HPLC (Ultimate 3000, Thermo scientific). The peptides were loaded in solvent A (0.1% formic acid in water) on a C18 reversed phase analytical column (Ionoptics, AUR3-15075C18-XT 15cm x 75um ID, 1.7um). The peptide mixture was resolved on a 5 to 28% linear gradient of solvent B (99.99% acetonitrile with 0.1% formic acid/) for 60 minutes followed by 15 minutes on a gradient of 28 to 95% and 15 minutes at 95% solvent B, at flow rates of 0.15 μl/min. Mass spectrometry was performed in a positive mode (m/z 300–1500, resolution 60, 000 for MS1 and 15, 000 for MS2, using repetitively full MS scan followed by high collision dissociation (HCD, at 27 normalized collision energy) of the 18 most dominant ions (>1 charges) selected from the first MS scan. The AGC settings were 3x106 for the full MS and 1x105 with maximum injection time of 60ms for the MS/MS scans. The intensity threshold for triggering MS/MS analysis was 1.7x104. A dynamic exclusion list was enabled with an exclusion duration of 20s.

The RAW files were converted to mzML files with MSConvert-GUI (68). The data was further analyzed by MSFragger with the LFQ-MBR workflow with default parameters (69). The downstream statistical analysis was performed on FragPipe-analyst (http://fragpipe-analyst.nesvilab.org/).

### Embryoid body differentiation

Non-directed differentiation was performed as in Shefa et al. (70). Cell aggregates were removed from an adherent culture of hESCs using Gentle Cell Dissociation Reagent (Stemcell technologies), and plated on Ultra-low attachment plates in DMEM/F12 supplemented with 5mM RI for 12-14 hours. Aggregates then transferred to differentiation medium (78% DMEM high glucose, 20% FBS, 0.1mM 2-mercaptoethanol, 1% NEAA, 3 mM L-glutamine, and Pen/Strep), medium replaced every 2 days. After 6 Days on Ultra-low attachment plates, aggregates were transferred to gelatin-coated plates at ∼10 EBs/cm2, and differentiation continued in the same medium.

### siRNA-mediated knockdown

hESCs in adherent culture were transfected with small interfering RNAs (siRNAs) using Lipofectamine RNAiMAX (Invitrogen) according to the manufacturer’s instructions. For each protein target, two pre-designed siRNAs targeting either RBM25 or CEBPZ (IDT) were used at a final concentration of 10nM. Transfections were performed twice, with a 24-hour interval between transfections to enhance KD efficiency. Cells were harvested 40 hours after the second transfection.

### Reverse transcription quantitative PCR

Total RNA was extracted from the cytoplasmic fraction obtained during the CUT&RUN procedure and purified using AMPure XP beads. cDNA was synthesized using the qScript cDNA Synthesis Kit (Quantabio) according to the manufacturer’s instructions. Quantitative PCR was performed using the PerfeCTa SYBR Green FastMix (Quantabio). Relative RNA expression levels were normalized to GAPDH.

### RNA sequencing

RNA sequencing for quantitative gene expression analysis was performed using the CEL-seq2 protocol (71), and libraries were sequenced on an Illumina NextSeq 2000 platform. Reads were trimmed using Cutadapt v3.5 and aligned to the reference genome using STAR v2.7.11b. Raw gene counts were generated using HTSeq-count v2.0.9 (72), and differential gene expression was analyzed using the DESeq2 R package (66).

For visualization of RNA expression in the UCSC Genome Browser, we performed full sequencing of hESCs RNA. Libraries were generated using NEBNext® rRNA depletion kit v2 (human/mouse/rat), followed by NEBNext® Ultra II RNA Library Prep Kit according to the manufacturer’s instructions. Libraries were sequenced on Illumina NEXTseq 2000 platform. The aligned BAM files from these libraries were used to display RNA-seq signal at selected genomic loci.

### ATAC-seq

ATAC-seq was performed as previously published (74). 50, 000 nuclei used for each sample, after transposase reaction DNA was purified and amplified for a total of 10 cycles. Paired end sequencing was performed on the Illumina NextSeq 2000 platform.

### Analysis of publicly available histone modification datasets

Publicly available histone modification ChIP-seq peak datasets were obtained from the ENCODE Project (GRCh38) (Supplementary Table S1). ENCODE-processed narrowPeak files were used for the analysis. The overlap between histone modification peaks and RBM25 CUT&RUN peaks located in promoter and intronic regions was assessed using BEDTools (53).

### Structural prediction of RBM25–nucleic acid interactions

Structural predictions of RBM25 interactions with RNA and DNA were performed using AlphaFold3 (75) with the full-length RBM25 protein. The DNA sequence GAGAGACTGCGCTTTCGGCCAATGAGTGCGTCGATTTCGAG was extracted from the CUT&RUN peaks including the CCAAT motif. The AlphaFold3-predicted complex was visualized and rendered using UCSF Chimera (76).

## Results

### RBM25 is identified as a candidate chromatin-associated RBP exhibiting extensive promoter co-occupancy with transcription factors

Recent studies indicate that RBPs can cooperate with TFs on chromatin, particularly at promoters, to regulate transcription (19, 36–38). To systematically identify chromatin-associated RBPs and their candidate TF partners in hESCs, we first performed promoter co-occupancy analysis using publicly available ENCODE ChIP-seq datasets from K562 cell line. After quality filtering (see Method), 127 TFs and 15 RBPs representing diverse protein families were retained for analysis (Supplementary Table S2). The TFs selected for analysis encompassed proteins from multiple DNA-binding domain (DBD) families, including C2H2 zinc finger (ZF), basic leucine zipper (bZIP), basic helix–loop–helix (bHLH), and others, classified according to the human TF atlas (77) (Figure S1A). The RBPs likewise represented diverse functional classes (19), including heterogeneous nuclear ribonucleoproteins (hnRNPs), serine/arginine-rich (SR) proteins, and RNA-binding motif (RBM)-containing proteins (Figure S1B).

Promoter co-occupancy was quantified for every TF–RBP pair by calculating the number of overlapping promoter peaks, followed by permutation testing to determine the significance of the observed overlap (Figure 1A and Supplementary Table S3). Among all RBPs analyzed, RBM25 exhibited a distinct promoter co-occupancy profile, showing significant overlap with multiple TFs, including NFY and SP TF families, as well as CEBPZ, NEUROD1, and several additional TFs. Based on its prominent promoter co-occupancy with multiple TFs in k562 cells and the expression levels of these factors in hESCs (Figure S1C), we chose to further explore its dual RNA and chromatin binding capacity in pluripotent cells.

**Figure 1:**
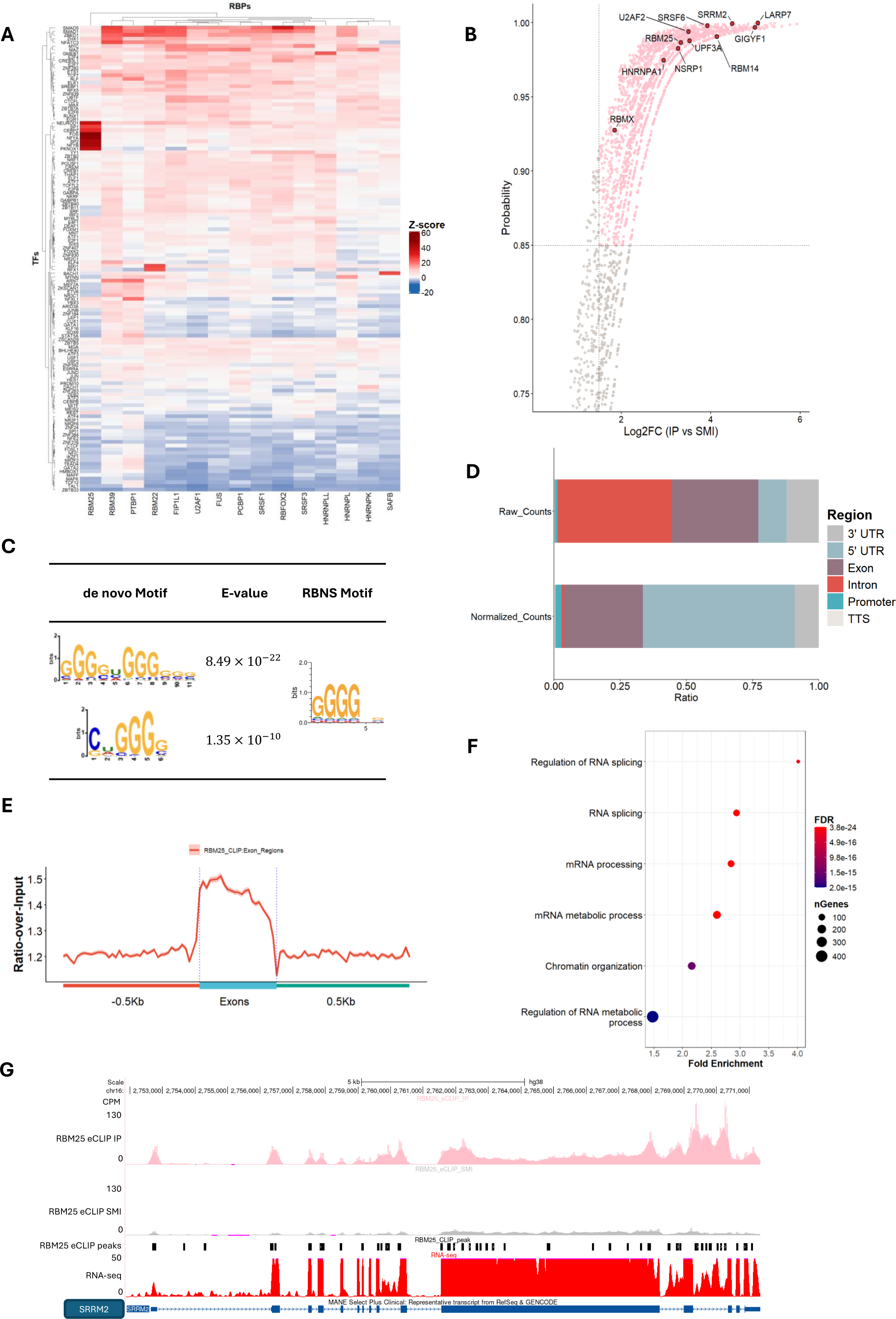
RBM25 binds RNA and functions as a splicing factor in hESCs. A: Heatmap showing the z-scores from a permutation test evaluating the enrichment of TFs and RBPs overlapped promoter-binding sites. B: Volcano plot of RBM25 eCLIP-enriched genes. Each dot represents a gene containing at least one cross-linked peak identified by PureCLIP. Log2 fold change (IP vs. SMI) and probability scores were calculated using NOISeq. Genes with a probability > 0.85 and log2FC > 1.5 were defined as RBM25-enriched targets and are highlighted in pink. Selected splicing factor genes are highlighted in red. C: De novo RNA-binding motif identified from RBM25 eCLIP peaks using XSTREME, together with the top-matched known RBP motifs and published RBNS motif. D: Distribution of RBM25 RNA-binding sites across transcript regions identified by eCLIP. Top, distribution based on raw peak counts. Bottom, distribution normalized by transcript region length. E: Metagene coverage profile of RBM25 binding across exons with 500-bp flanking regions on both sides. F: Gene Ontology enrichment analysis (for biological processes) of RBM25 RNA targets. G: UCSC Genome Browser view of the SRRM2 locus displaying RBM25 CLIP IP and SMI bigWig signal tracks, significant cross-linked peaks, and RNA-seq signal.

### RBM25 RNA binding pattern in hESCs supports its canonical role in splicing regulation

To systematically identify RBM25 binding targets and functions at the RNA level in hESCs, we performed enhanced crosslinking and immunoprecipitation sequencing (eCLIP-seq) (47). Significant RBM25-binding sites were identified using PureCLIP followed by NOISeq comparison between immunoprecipitated (IP) and size-matched input (SMI) libraries (Figure 1B). In total, we identified 6, 936 significant RBM25-binding peaks corresponding to 1, 592 genes (Supplementary Table S4). Notably, RBM25 bound numerous transcripts encoding splicing regulators, as expected based on the prevalence of cross-regulatory networks within the splicing machinery. Among the RBM25 eCLIP targets we found SRRM2, U2AF2, RBM14, and SRSF6 (78). De novo motif analysis identified G-rich sequence motifs as the predominant RBM25-binding motif (Figure 1C), consistent with its RNA-binding preferences previously determined by in vitro RNA Bind-n-Seq (RBNS) assay (79).

To characterize the distribution of RBM25 binding sites, eCLIP peaks were annotated to transcript features. Based on absolute peak counts, we found that RBM25-binding sites are predominantly located within intronic and exonic regions (Figure 1D, top). Because transcript features differ substantially in length, peak densities were normalized to feature length. After normalization, RBM25 binding was found to be most strongly enriched in 5′ untranslated regions (5′UTRs) and exons (Figure 1D, bottom). Metagene analysis further revealed that RBM25 binding is concentrated around exon boundaries, with prominent enrichment near the 3′ splice site (Figure 1E), supporting its well-established function as a splicing factor. Functional enrichment analysis further demonstrated that RBM25 target genes are significantly associated with RNA splicing and mRNA processing pathways (Figure 1F). Among the most enriched targets was the mRNA encoding SRRM2 protein (Figure 1B and 1G). SRRM2 is an essential spliceosome component that has been implicated in stem cell development and human disease (80–83). Visualization of the SRRM2 locus in the UCSC Genome Browser shows that RBM25-binding sites are predominantly located within the gene’s exons, with additional binding sites detected in introns and untranslated regions (Figure 1G). Together, our results demonstrate that RBM25 preferentially binds exonic regions of splicing-related transcripts and recognizes a G-rich RNA motif in hESCs, consistent with its canonical splicing-associated function.

### RBM25 exhibits distinct promoter and intronic chromatin-binding landscapes in hESCs

To define the chromatin-binding landscape of RBM25 in hESCs, we performed CUT&RUN. Comparison with IgG control libraries identified 13, 406 high-confidence RBM25 chromatin-binding peaks associated with 9, 121 genes (Supplementary Table S5). RBM25 peaks were found to be mainly located in intronic regions, followed by promoters (Figure 2A). As several splicing factors have been reported to bind both promoters and introns of the same genes, supporting co-transcriptional splicing (28, 35), we examined whether RBM25 showed a similar pattern in hESCs. However, only a small subset of genes contained RBM25 peaks in both promoter and intronic regions (Figure 2B). These results indicate that promoter- and intron-associated RBM25 binding predominantly occurs at distinct gene sets.

**Figure 2:**
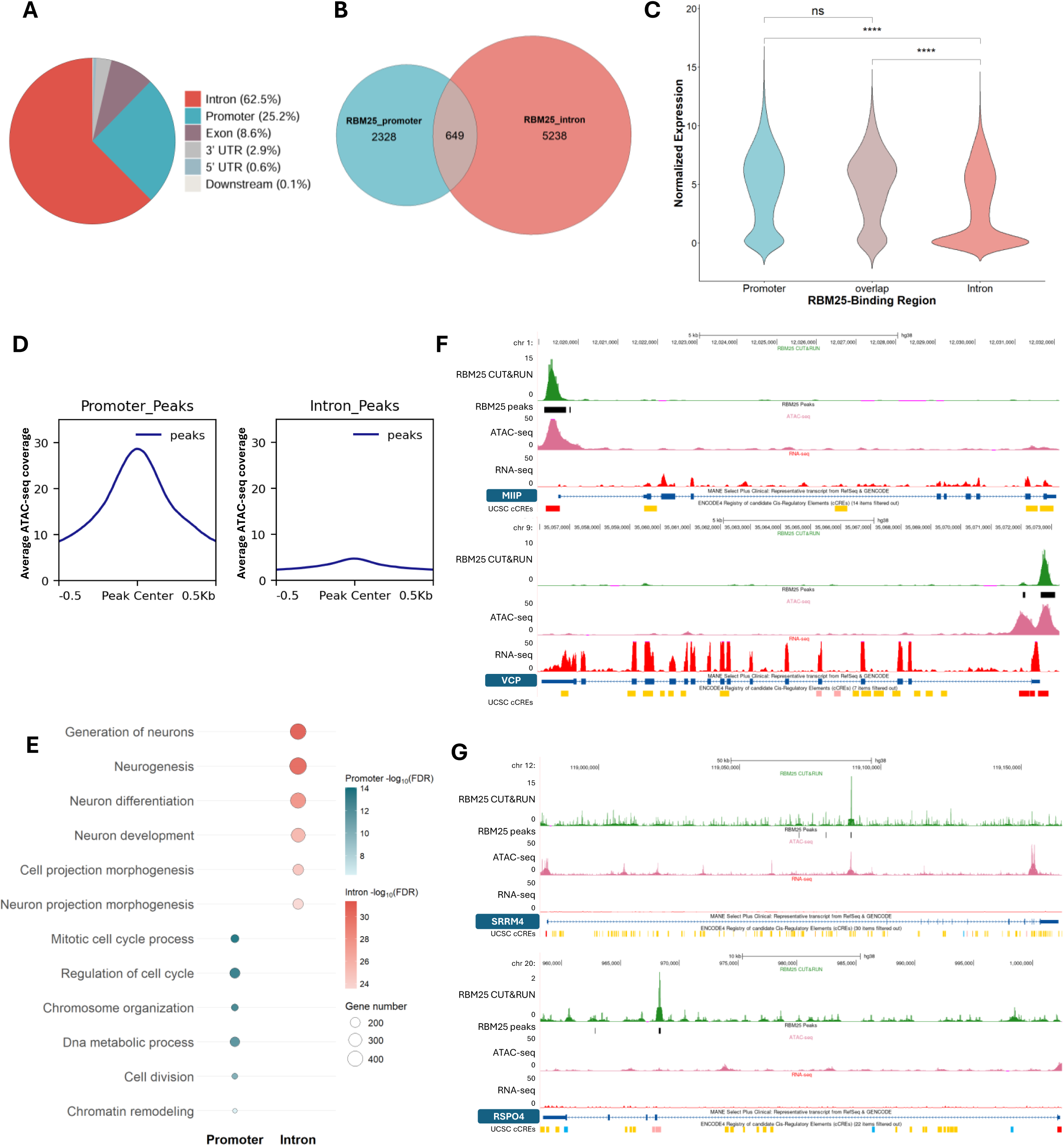
RBM25 occupies distinct promoter and intronic chromatin regions in hESCs. A. Genomic distribution of RBM25 chromatin-binding peaks identified by CUT&RUN. B. Overlap between genes with promoter-localized RBM25 peaks and genes with intronic RBM25 peaks. C. Violin plot comparing normalized expression levels of genes with promoter-only, intron-only, or both promoter and intronic RBM25 binding. Statistical significance was assessed by t-test. D. ATAC-seq signal over RBM25 CUT&RUN peaks. Left, promoter-localized peaks. Right, intronic peaks. E. Gene Ontology enrichment analysis (for biological processes) of genes with promoter-localized or intronic RBM25 binding. F. UCSC Genome Browser views showing RBM25 CUT&RUN signal, RBM25 peaks, ATAC-seq signal, and RNA-seq signal at representative promoter-bound loci. Top, MIIP. Bottom, VCP. G. UCSC Genome Browser views showing RBM25 CUT&RUN signal, RBM25 peaks, ATAC-seq signal, and RNA-seq signal at representative intron-bound loci. Top, RSPO4. Bottom, PLXNA4.

We next integrated RNA-seq data to compare transcriptional activity across these target classes. Promoter-bound genes were found to be generally highly expressed, whereas intron-bound genes are predominantly lowly expressed or transcriptionally silent. Genes with both promoter and intronic RBM25 binding showed expression levels comparable to promoter-bound genes (Figure 2C). Because protein-coding genes and non-coding RNAs have distinct transcriptome-wide expression distributions (84), we further stratified the analysis by gene biotype. Most promoter-bound and intron-bound genes were classified as protein-coding (72% and 60%, respectively), followed by long non-coding RNAs (27% and 37% for promoter-bound and intron-bound, respectively) (Figure S2A). Within both biotype classes, promoter-bound genes consistently showed higher expression than intron-bound genes. As expected from the distinct expression distributions of different gene biotypes (84), protein-coding genes showed substantially higher expression levels than non-coding genes across both promoter-bound and intron-bound groups (Figure S2B). In promoter-bound genes we also noticed a strong ATAC-seq signal around the Transcription Start Sites (TSS), whereas the TSSs of intron-bound genes were detected as largely inaccessible (Figure S2C). The ATAC-seq data together with the RNA-seq data strongly indicate that promoter-bound RBM25 targets define an actively transcribed and accessible gene set, whereas intron-bound targets are generally inactive. We then asked whether the local chromatin environment differed between the promoter and intronic RBM25 peaks themselves. Expectedly, promoter-localized RBM25 peaks were characterized by strong ATAC-seq signal, consistent with an open chromatin state (Figure 2D, left). In contrast, intronic RBM25 peaks showed substantially lower ATAC-seq signal (Figure 2D, right). Gene Ontology analysis further separated the two target classes. Promoter-bound genes were found to be enriched for cell cycle-related processes, whereas intron-bound genes were enriched for neuronal development-related genes (Figure 2E).

Among the promoter-bound targets of RBM25 were included MIIP and VCP, which have reported roles in proliferation, DNA repair, and cell cycle regulation (85–89). At both loci, RBM25 peaks were detected within accessible promoters of actively expressed genes (Figure 2F). Among the intron-bound targets we observed PLXNA4 and RSPO4, both which are transcriptionally silent in hESCs. PLXNA4 is known for its role in neuronal differentiation and axon guidance (90) and RSPO4, a protein coding gene involved in nail development (91). Interestingly, the intronic RBM25 peaks within these genes differed in their chromatin accessibility. While the PLXNA4 peak was found to overlap an open chromatin region, the RSPO4 peak was located with a relatively inaccessible region (Figure 2G). Genes with both promoter and intronic RBM25 binding included NOVA2 and CYB5B, which displayed distinct local chromatin features (Figure S2D).

To further characterize the chromatin environment associated with RBM25 binding, we integrated RBM25 CUT&RUN peaks with ATAC-seq and ENCODE histone modification datasets (Supplementary Table S1). Most RBM25 promoter-binding peaks were found to overlap active chromatin features. Specifically, 82% overlapped H3K4me3, 79% overlapped ATAC-seq peaks, and 74% overlapped both features (Figure S2E). As H3K4me3 chromatin mark is generally associated with active transcription initiation (92, 93), these results support the preference of RBM25 to bind within active promoters. In contrast, intronic RBM25 peaks showed much lower overlap with active chromatin features. Only 12% overlapped ATAC-seq peaks, and 11% overlapped H3K4me3 peaks (Figure S2F). Because intronic H3K4me3 has been linked to co-transcriptional pre-mRNA splicing (94), we next tested whether stronger intronic RBM25 peaks are preferentially associated with local chromatin accessibility and H3K4me3 enrichment. Minimum hypergeometric (mHG) enrichment analysis showed a positive association between intronic RBM25 peak strength and overlap with both ATAC-seq and H3K4me3 peaks (P = 1.13 ×10⁻⁸ and 1.11 ×10⁻⁸, respectively; Figure S2G, H). Approximately 70% of H3K4me3-positive intron-bound genes are transcriptionally active, compared with approximately 50% of genes where RBM25 did not overlap with H3K4me3 marks (Figure S2I). Thus, although most intronic RBM25 binding occurs at inaccessible or inactive regions, a subset is associated with transcriptionally permissive chromatin. Overall, our findings identify two major classes of RBM25 chromatin targets in hESCs. Promoter-associated RBM25 binding which occur mainly at accessible, actively expressed cell cycle genes. Intron-associated binding that are enriched at neuronal genes and are largely transcriptionally silent, with a smaller subset marked by active chromatin features. To further investigate the potential of RBM25 to interact directly with DNA, we modeled the interaction between the full protein and a selected DNA sequence extracted from the CUT&RUN top peaks using AlphaFold 3. As shown in Supplementary Figure S2J, the predicted structure of the RBM25-DNA complex suggests a potential DNA-binding interface primarily involving the RRM domain of RBM25. Note that the full protein structure (not shown) is predicted as largely disordered. Interestingly, while in previous studies it has been proposed that the PWI domain of RBM25 can interact with both single- and double-stranded nucleic acids (44), based on AlphaFlod predictions the PWI domain has no shared interface with the DNA

### RBM25 RNA- and chromatin-associated targets in hESCs are functionally distinct

Having characterized the RNA- and chromatin-binding landscapes of RBM25, we then asked whether these activities converged on the same target genes. Several DNA- and RNA-binding proteins have been reported to bind the same genes at both the chromatin and RNA levels, thereby coordinating transcriptional and post-transcriptional regulation of the same genes (28, 35, 36, 95). We therefore compared RBM25 targets identified by eCLIP and CUT&RUN. Among the 1, 592 RNA-bound genes identified by eCLIP and the 9, 121 chromatin-bound genes identified by CUT&RUN, we found 621 genes were shared, higher than expected by chance (Figure S3A). Nevertheless, the shared genes represented only 39.0% of the eCLIP targets and 6.8% of the CUT&RUN targets, indicating that the majority of RBM25 targets identified by the two assays were distinct (Figure S3A). Metagene analysis showed that RBM25 RNA-binding sites are broadly distributed across gene bodies. Consistent with the promoter-associated chromatin occupancy of RBM25 reported in other cell types (19), RBM25 chromatin-binding sites in hESCs showed pronounced enrichment around TSSs (Figure S3B).

Gene Ontology analysis further supported the distinction between the RNA- and chromatin-associated target sets of RBM25. While eCLIP-specific targets were enriched for RNA splicing and RNA processing terms, chromatin-bound targets showed strong enrichment for neuronal development and cell cycle-related processes. Some GO terms were shared between the RNA- and chromatin-associated target sets, which may reflect the contribution of genes bound by RBM25 at both the RNA and chromatin levels. Notably, enrichment for neuronal development and cell cycle-related processes among these shared targets was substantially less significant than that observed for CUT&RUN-specific targets (Figure S3C). Moreover, there were no positional preferences for these sites (Figure S3E). Overall, the data indicates that RBM25 functions as both an RNA- and chromatin-binding protein in hESCs, with these two activities targeting largely distinct gene sets involved in different biological processes.

### CEBPZ is identified as a candidate chromatin-associated partner of RBM25

After establishing that RBM25 targets distinct genes at the RNA and chromatin levels, we sought to identify TFs that may associate with RBM25 on chromatin. Previous studies have shown that chromatin-associated RBPs can co-occupy regulatory regions with transcription factors, and that such TF-RBP co-binding is often accompanied by shared or enriched DNA sequence motifs (36, 38). De novo motif analysis of RBM25 CUT&RUN peaks identified three highly enriched sequence motifs: a CCAAT motif, known to be recognized by major transcription factor families NFY (96); a GC-box motif matching the binding motifs of SP1 and SP2 TFs, and a CTCF-like motif (Figure 3A). Notably, these TFs were also predicted as the top RBM25 promoter co-occupancy partners in the ENCODE-based analysis (Figure 1A), providing independent support for their association with RBM25-bound chromatin regions. We next examined motif enrichment separately in promoter- and intron-associated RBM25 peaks. The CCAAT and GC-box motifs were preferentially enriched in promoter-associated RBM25 peaks, whereas the CTCF-like motif was predominantly enriched in intronic peaks (Figure S4A). Consistent with this promoter-biased enrichment, approximately half of all CCAAT-containing RBM25 peaks were found in promoter regions, followed by introns (Figure S4B). Moreover, among the RBM25 binding genes, those containing a CCAAT motif exhibited significantly higher expression levels than those lacking the motif (Figure 3B). These results suggest that CCAAT-containing promoter elements define an active subset of RBM25-bound chromatin regions. Consistent with these results, the Alphafold3 model of protein-DNA complex predicts that the loop of the RRM faces the double stranded DNA at the CCAAT motif, particularly at the AAT nucleotides (Figure S3C).

**Figure 3:**
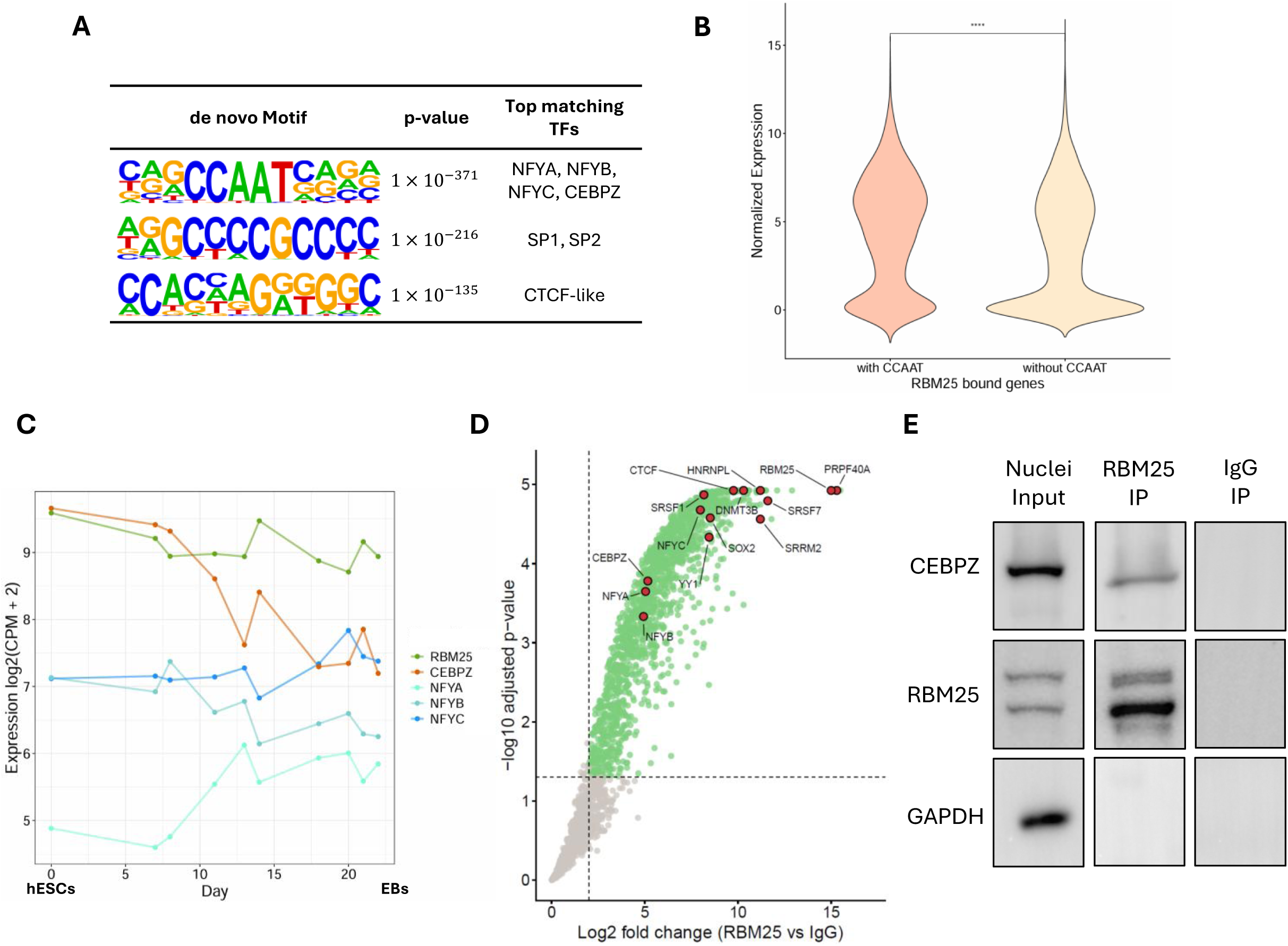
CEBPZ is identified as a chromatin-associated interaction partner of RBM25. A. De novo sequence motifs enriched within RBM25 CUT&RUN peaks, identified using HOMER, together with the top-matched known TF motifs. B. Violin plot comparing normalized expression levels of RBM25-bound genes with or without a CCAAT motif within the corresponding RBM25 peak. Statistical significance was assessed by t-test. C. Expression dynamics of RBM25, CEBPZ, and NFY family members during differentiation from hESCs (day 0) to embryoid bodies (day 21). Correlations with RBM25 expression were calculated using Spearman’s correlation coefficient: RBM25-CEBPZ: 0.77, RBM25-NFYA: -0.73, RBM25-NFYB: 0.12, RBM25-NFYC=-0.64. D. Co-immunoprecipitation followed by mass spectrometry comparing RBM25 immunoprecipitate with IgG controls. Each point represents a detected protein. Proteins with adjusted P < 0.05 and log2 fold change > 2 were defined as significantly enriched and are highlighted in green. Selected proteins are highlighted in red. E. Western blot validation of RBM25 co-immunoprecipitation using nuclear lysates. GAPDH is shown as a control.

Given the strong enrichment of the CCAAT motif at promoter-associated RBM25 peaks, we asked which CCAAT-binding TFs was most closely associated with RBM25 expression in hESCs. Analysis of RNA-seq data across differentiation from hESCs to embryoid bodies revealed that CEBPZ was the only CCAAT-binding candidate that showed a significant positive correlation with RBM25 (Figure 3D). Moreover, CEBPZ was expressed at significantly higher levels compared to the NFY family members in hESCs (Figure 3D and Figure S1C). Together, these data strengthen the notion that RBM25 chromatin binding could be associated with CEBPZ.

To test whether RBM25 and CEBPZ interact physically, we performed RBM25 co-immunoprecipitation followed by mass spectrometry. This analysis identified 1, 674 proteins significantly enriched in RBM25 IP compared with IgG controls, including CEBPZ which ranked 982 of 1674 enriched genes by the Log2FC. (Figure 3E and Supplementary Table S6). Consistent with the RNA-processing role of RBM25 in hESCs, many of the other top ranked proteins are known splicing factors. The association between RBM25 and CEBPZ was further validated by anti-RBM25 co-IP from nuclear lysates and reciprocal co-immunoprecipitation from whole-cell lysates (Figure 3F and Figure S4D).

### RBM25 and CEBPZ exhibit similar DNA-binding patterns

Following our observation that RBM25 physically interacts with CEBPZ in hESCs, we sought to characterize the chromatin-binding properties of CEBPZ in this stem cell model. To define the chromatin-binding landscape of CEBPZ in hESCs, we performed CUT&RUN. Comparison with IgG control libraries identified 9, 890 high-confidence CEBPZ chromatin-binding peaks associated with 7, 004 genes (Supplementary Table S7). De-novo motif analysis identified GC-box and CCAAT motifs as the two most enriched motifs within CEBPZ peaks (Figure S5A), similar with the motif profile observed for RBM25 (Figure 3A). Genome-wide annotation showed that CEBPZ peaks were mainly located in intronic regions, followed by promoter regions (Figure S5B). As observed for RBM25, genes with promoter-associated CEBPZ binding are expressed at significantly higher levels than genes possessing intronic CEBPZ binding sites (Figure S5C). Here again, GO analysis indicated an enrichment for cell cycle-related processes, whereas intron-bound genes were found to be enriched for neuronal development related genes (Figure S5D). Metagene analysis showed broadly similar gene-body distributions for RBM25 and CEBPZ. However, RBM25 showed stronger enrichment around TSSs, whereas CEBPZ displayed broader occupancy across gene bodies (Figure 4A). Together, these findings indicate that RBM25 and CEBPZ share several chromatin-binding features.

**Figure 4:**
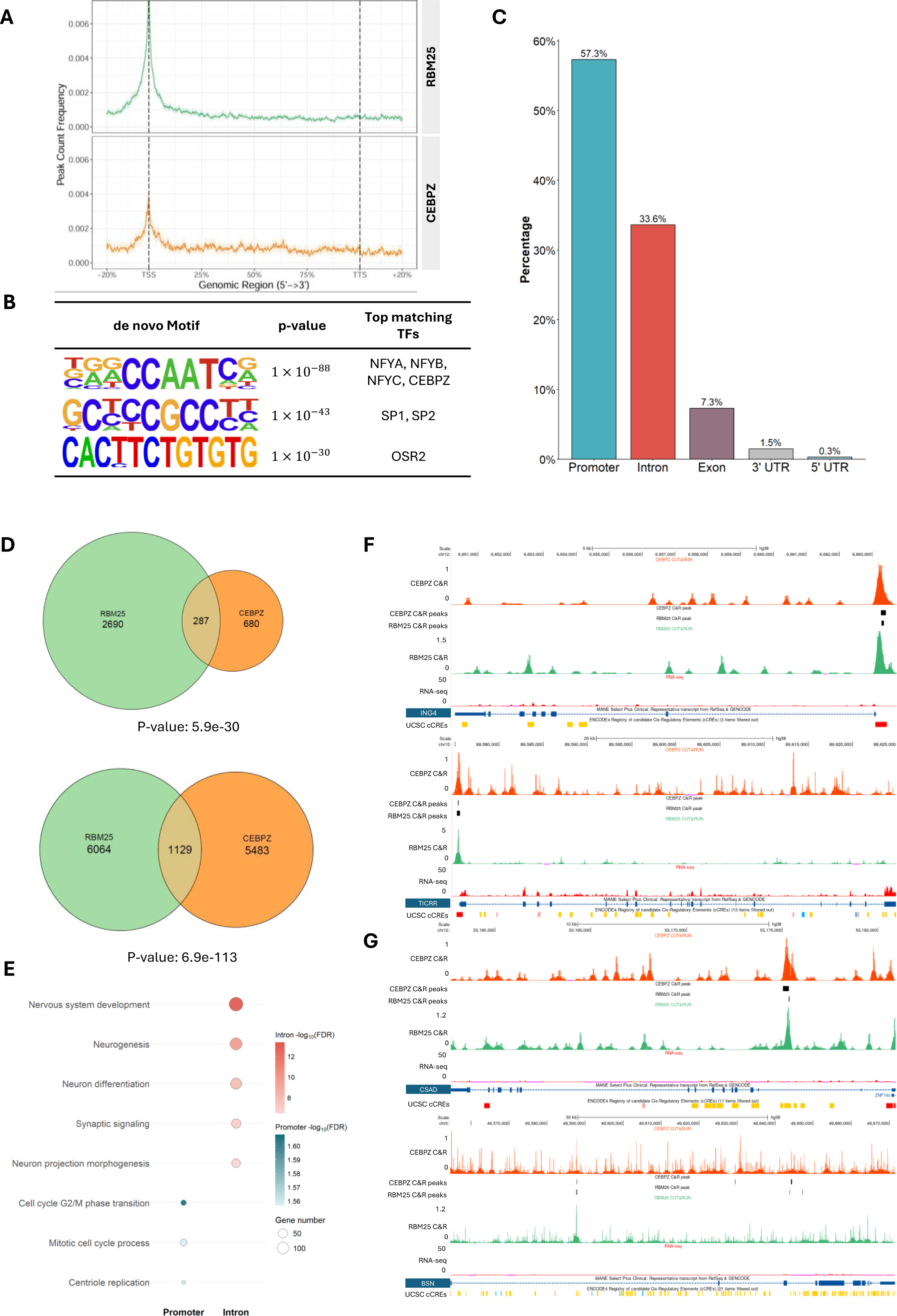
RBM25 and CEBPZ exhibit shared DNA-binding properties. A: Metagene coverage plots showing RBM25 and CEBPZ binding distributions across gene bodies based on CUT&RUN data. B: De novo DNA-binding motifs enriched within CUT&RUN peaks bound by both RBM25 and CEBPZ identified using HOMER, together with their top-matched TF motifs. C: Distribution of peaks bound by both RBM25 and CEBPZ across genomic regions. D: Overlap analysis of RBM25 and CEBPZ binding within specific genomic regions. Top: Promoter regions. Bottom: Intronic regions. Statistical significance was assessed using a chi-square test. E: Gene Ontology enrichment analysis (for biological processes) of RBM25&CEBPZ co-binding genes bound in promoter and intronic regions. F: Representative examples of genes bound by both RBM25 and CEBPZ at promoter regions, shown in UCSC genome browser view. Top: ING4. Bottom: TICRR. Gene expression levels are indicated by RNA-seq signal. G: Representative examples of genes bound by both RBM25 and CEBPZ within intronic regions, shown in UCSC genome browser view. Top: CSAD. Bottom: BSN. Gene expression levels are indicated by RNA-seq signal.

We next examined genomic regions co-occupied by RBM25 and CEBPZ. De novo motif analysis of shared CUT&RUN peaks again identified the CCAAT motif as the most enriched motif (Figure 4B), consistent with the individual motif analyses of RBM25 and CEBPZ. Most CCAAT-containing shared peaks were found within promoter regions (Figure 4C). Genes associated with shared CCAAT-containing peaks were also expressed at significantly higher levels than genes associated with shared peaks lacking this motif (Figure S6A). These results indicate that CCAAT-containing promoter elements mark the active subsets of RBM25 and CEBPZ co-occupied regions. In parallel, analysis of RBM25 and CEBPZ co-occupancy across genomic regions revealed that co-occupied peaks were more prevalent within intronic regions than promoter regions. (Figure S6B). Chi-square analysis showed that RBM25 and CEBPZ co-occupancy were significantly enriched in both promoter and intronic regions compared to random (Figure 4D). Further analysis showed that intronic co-occupancy was the most frequent pattern, followed by promoter co-occupancy (Figure S6C). GO analysis revealed that promoters of genes bound by both RBM25 and CEBPZ were enriched for cell cycle-related processes, whereas genes bound by both factors at the intron region are associated with neuronal development (Figure 4E). Examples of promoter bound genes involved in cell proliferation, apoptosis, and DNA replication (ING4 and TICRR(97, 98)) and others bound with introns and associated with neuronal differentiation and synaptic activity (CSAD and BSN(99, 100)) are shown in Figure 4F and 4G, respectively. This data is consistent with the assumption that RBM25 acts in conjunction with CEBPZ to regulate the expression of genes in hESCs. While promoter binding seems to be associated with positive regulation of cell cycle and proliferation genes while intron binding is associated with negative regulation of neuronal related genes.

### RBM25 promotes CEBPZ occupancy at CCAAT-containing promoters of cell cycle genes

To investigate the functional relationship between RBM25 and CEBPZ, we performed siRNA-mediated knockdown (KD) of each factor followed by RNA-seq. Efficient depletion was confirmed at both the RNA and protein levels (Figures 5A and S7A). CEBPZ depletion altered the expression of 99 genes, comprising 91 upregulated and 8 downregulated genes. By comparison, RBM25 depletion altered 65 genes, comprising 27 upregulated and 38 downregulated genes (Figures S7B, C and Supplementary Table S8). Gene expression changes induced by the two knockdowns showed a positive correlation (Figure S7D), consistent with the model suggesting cooperation between the two factors in regulating unique subset of genes in hESCs. Notably, KD of each factor did not result in a significant change in the expression of the other (Supplementary Table S8).

**Figure 5:**
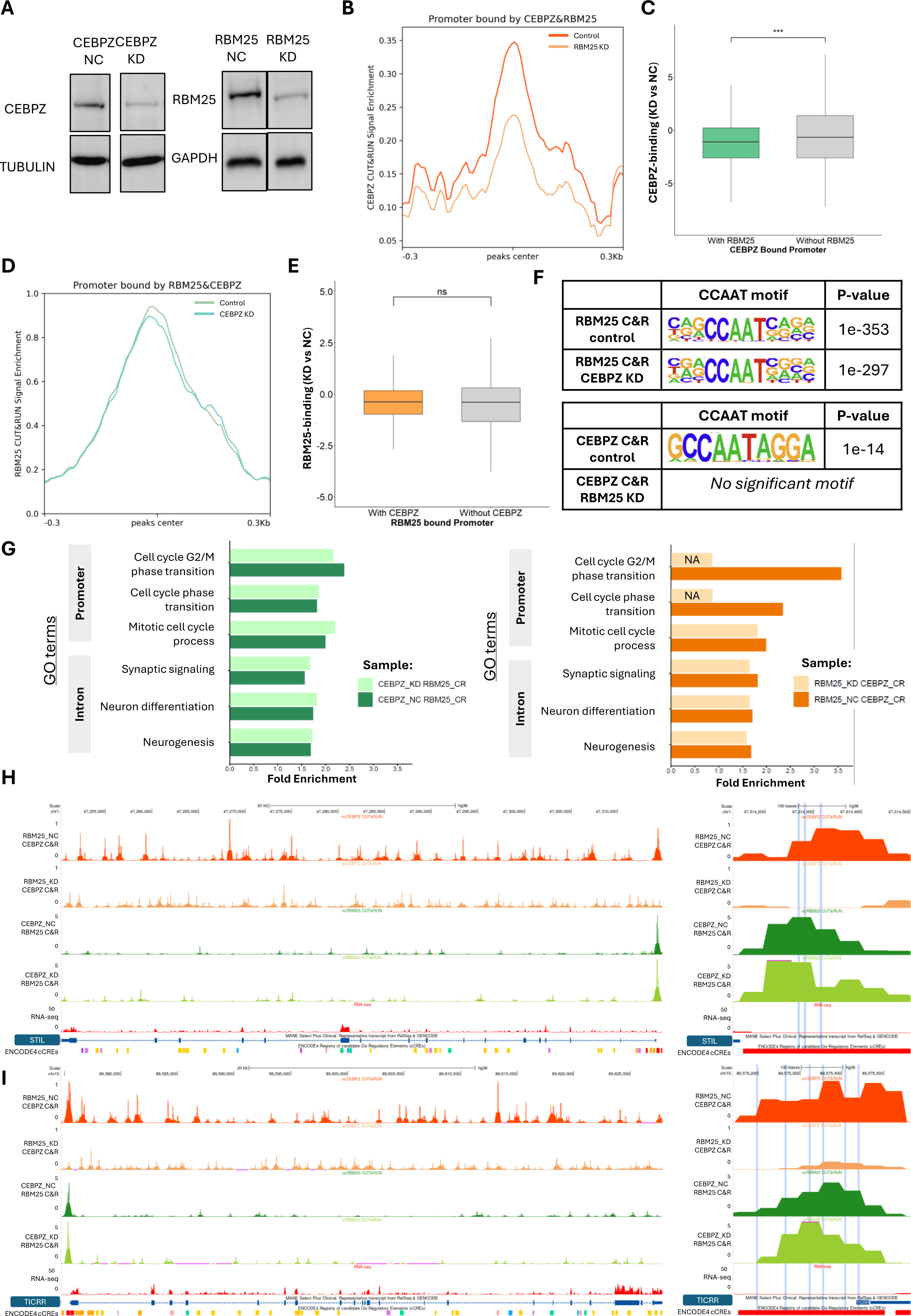
RBM25 is required for CEBPZ occupancy at promoters of cell cycle genes. A. Western blot analysis of RBM25 and CEBPZ protein levels following individual siRNA-mediated knockdown (KD) and the corresponding control treatments. B. Metagene profiles of CEBPZ CUT&RUN signal across promoter regions bound by both CEBPPZ and RBM25 in control and RBM25-depleted cells. C. Distribution of log2FC in CEBPZ CUT&RUN signal following RBM25 KD at promoter regions bound by both factors and CEBPZ-specific promoter regions. Statistical significance was assessed using Student’s t-test. D. Metagene profiles of RBM25 CUT&RUN signal across promoter regions bound by both CEBPPZ and RBM25 in control and CEBPZ-depleted cells. E. Distribution of log2FC in RBM25 CUT&RUN signal following CEBPZ KD at promoter regions bound by both factors and RBM25-specific promoter regions. Statistical significance was assessed using Student’s t-test. F. De novo motif analysis of RBM25 and CEBPZ CUT&RUN peaks before and after reciprocal KD. Left, motifs enriched within RBM25 peaks in control and CEBPZ-depleted cells. Right, motifs enriched within CEBPZ peaks in control and RBM25-depleted cells. G. Gene Ontology enrichment analysis (for biological processes) of genes associated with RBM25 or CEBPZ CUT&RUN peaks before and after reciprocal KD. Left, RBM25-associated genes in control and CEBPZ-depleted cells. Right, CEBPZ-associated genes in control and RBM25-depleted cells. H. UCSC Genome Browser views of RBM25 and CEBPZ occupancy at the STIL locus before and after reciprocal KD. Left, genomic view of the locus. Right, enlarged view of the promoter region. RBM25 and CEBPZ CUT&RUN and RNA-seq signals are shown. I. UCSC Genome Browser views of RBM25 and CEBPZ occupancy at the TICRR locus before and after reciprocal KD. Left, genomic view of the locus. Right, enlarged view of the promoter region. RBM25 and CEBPZ CUT&RUN and RNA-seq signals are shown.

To determine whether either factor depended on the other for chromatin occupancy, we performed reciprocal CUT&RUN following depletion of the interacting factor. We compared normalized CUT&RUN signals and log2 fold changes at regions bound by both factors and factor-specific promoter and intronic peaks. RBM25 depletion significantly reduced CEBPZ occupancy specifically at promoter peaks that were also found to bind RBM25 (Figure 5B&C, S8A). In contrast, CEBPZ depletion did not result in a comparable reduction in RBM25 occupancy at shared promoter peaks (Figure 5D&E, S8B). At intronic regions, depletion of either RBM25 or CEBPZ did not significantly reduce occupancy of the other factor (Figure S8C–F). These results reveal a unidirectional dependency in which RBM25 supports CEBPZ occupancy at promoter regions, whereas no reciprocal dependency was detected, or any dependency at intronic regions.

We next examined whether this dependency was associated with changes in the motif composition. Notably, the CCAAT motif remained strongly enriched among RBM25 peaks following CEBPZ depletion. In contrast, CCAAT motif enrichment was markedly reduced among CEBPZ peaks following RBM25 depletion (Figure 5F). This selective reduction was consistent with preferential loss of CEBPZ occupancy at CCAAT-containing regions. GO analysis further showed that genes associated with CEBPZ peaks were no longer enriched for cell cycle phase transition-related processes following RBM25 depletion. Enrichment for most other biological processes remained largely unchanged (Figure 5G). This promoter-specific dependency was evident at TICRR and STIL, two cell cycle regulators found to be bound by both RBM25 and CEBPZ in hESCs (Figure 5H, I). TICRR is required for DNA replication and cell cycle checkpoint regulation (98, 101), whereas STIL contributes to centriole biogenesis and cell cycle progression (102, 103). At both loci, RBM25 occupancy was maintained following CEBPZ depletion, whereas CEBPZ occupancy was markedly reduced following RBM25 depletion. Both promoter regions contained multiple CCAAT motifs, further linking RBM25-dependent CEBPZ occupancy to CCAAT-containing promoters of cell cycle genes. Together, these results show that RBM25 is required for CEBPZ binding at promoters that are targeted by both factors, specifically CCAAT-containing promoters of cell cycle genes. In contrast, reciprocal dependence was not detected at the intronic regions associated with neuronal genes, which were also found to be targeted by both factors. Thus, although at the chromatin level, RBM25 and CEBPZ co-occupy both promoter and intronic regions, their functional interdependence is predominantly promoter-associated in hESCs.

## Discussion

In this study we investigate the role of RBM25 as a dual-function DRBP in hESCs. We show that at the RNA level, RBM25 preferentially binds transcripts enriched for RNA processing and splicing factors, consistent with its known role as an RNA and splicing regulator. At the chromatin level, however, RBM25 exhibits context-dependent binding associated with distinct functional outcomes at promoters and introns. At promoter regions, RBM25 binds to actively expressed genes enriched for cell cycle–related functions and acts as a transcriptional co-factor, facilitating the recruitment of the TF CEBPZ to its well established CCAAT motifs. In contrast, within the intronic regions, RBM25 binds predominantly to transcriptionally inactive genes enriched for neuronal functions. Many of the genes bound by RBM25 within introns were also found to be bound by CEBPZ, suggesting a potential functional cooperation between the two factors in the context of chromatin binding. These results support a model in which RBM25 integrates multiple layers of gene regulation in hESCs: coordinating transcriptional control through chromatin binding and TF recruitment at promoters, while simultaneously regulating RNA processing. We propose that through these dual activities, and specifically its unique chromatin binding preferences, RBM25 may contribute to human pluripotency.

RBM25 has been extensively characterized as a splicing factor in multiple cell types, where it regulates alternative splicing through interactions with pre-mRNAs and the spliceosome (39–43). Although we did not directly examine RBM25-dependent splicing events, our findings support that RBM25 retains its canonical RNA-associated function in hESCs. Using eCLIP-seq, we provide, to our knowledge, the first transcriptome-wide map of RBM25 RNA-binding sites generated by a high-throughput assay. RBM25 directly binds transcripts enriched for RNA processing and pre-mRNA splicing functions. Consistent with these findings, co-immunoprecipitation coupled with mass spectrometry identified numerous RBM25-interacting proteins involved in pre-mRNA splicing, including members of the SR and hnRNP families, in agreement with previous reports that RBM25 associates with spliceosome-related ribonucleoprotein complexes (39). Together, our RNA-binding and proteomic analyses reinforce the canonical RNA-associated role of RBM25 in the hESCs. Future studies integrating high-depth RNA-seq with splicing-aware analyses will be required to identify direct splicing targets and define the broad contribution of RBM25 to alternative splicing in hESCs.

An increasing number of RBPs have been shown to associate with chromatin in addition to their classical RNA-binding activities, thereby functioning as DRBPs that connect transcriptional and post-transcriptional regulation (35, 104). Previous studies identified RBM25 as a chromatin-associated protein and demonstrated that it can cooperate with transcription factors in a cell-type-dependent manner, including by promoting YY1 occupancy at active promoters in HepG2 cells (19). In a later study, deletion of RBM25 affected dramatically the gene expression in mESCs (45). However, in our study, there was only a small number of differentially expressed genes, with no clear functional enrichment, upon KD of RBM25. The relatively modest transcriptional effect may partly reflect incomplete RBM25 depletion. Our CUT&RUN analysis extends previous observations by defining the chromatin-binding landscape of RBM25 in hESCs and revealing two distinct and novel modes of occupancy. Promoter-associated binding occurred predominantly at accessible, actively transcribed genes enriched for cell cycle-related functions, whereas intronic binding was primarily associated with transcriptionally inactive neuronal genes. Promoter-bound RBM25 extensively overlapped open chromatin and active histone marks, whereas most intronic peaks occurred in relatively inaccessible chromatin. Although a subset of intronic peaks overlapped H3K4me3-marked regions potentially associated with transcription-coupled RNA processing (105, 106), this subset may represent sites where chromatin-associated RBM25 participates in active transcriptional regulation. However, such a mechanism is unlikely to explain the majority of intronic RBM25 binding events, particularly those associated with transcriptionally inactive neuronal genes. Thus, distinct molecular mechanisms may recruit or retain RBM25 at promoter and intronic chromatin. Comparison of the eCLIP and CUT&RUN datasets revealed limited overlap between the RNA- and chromatin-associated targets of RBM25, which also showed distinct genomic distributions and functional enrichments. These findings suggest that the RNA- and chromatin-associated activities of RBM25 largely represent distinct regulatory programs, potentially determined by different molecular partners.

Interactions between RBPs and transcription factors have emerged as an important mechanism through which chromatin-associated RBPs can influence transcription (19, 36–38). Our findings identify RBM25 as a chromatin-associated partner of the TF CEBPZ in hESCs, supported by their physical association and extensive genomic co-occupancy in these cells. Importantly, this association was not reported before in other cell lines and is likely context-dependent. At shared promoters, RBM25 depletion reduced CEBPZ occupancy, whereas CEBPZ depletion had little effect on RBM25 binding, suggesting that RBM25 facilitates CEBPZ association with CCAAT-containing promoters. In contrast, their binding at intronic regions was largely independent. Together, these findings suggest a role for RBM25 in promoting promoter-specific occupancy of CEBPZ in pluripotent cells. Notably, although gene expression changes following RBM25 and CEBPZ depletion were positively correlated, CEBPZ depletion resulted in a distinct and larger set of differentially expressed genes. Only a small subset of these genes was directly bound by CEBPZ, suggesting that many of the observed transcriptional changes may reflect indirect downstream effects. Moreover, the relatively modest transcriptional responses to both KDs may partly result from incomplete depletion, with residual protein potentially sufficient to maintain some of their regulatory activities. More complete depletion, such as through genetic knockout, will be required to further define the transcriptional functions of RBM25 and CEBPZ in hESCs.

While RBM25 chromatin occupancy shows a clear preference for CCAAT-containing regions, the mechanism underlying this specificity remains unclear. Although the PWI domain of RBM25 has been reported to bind double-stranded DNA (44), there is currently no evidence linking this domain to sequence-specific recognition. Our structural prediction instead suggests that the RRM domain, rather than the PWI domain, is involved in DNA interaction. Moreover, CUT&RUN cannot distinguish direct DNA binding from indirect chromatin association mediated by protein–protein interactions (61), leaving open the possibility that RBM25 is recruited through other transcription factors. In addition to CEBPZ, NFY components may contribute to CCAAT recognition, as NFYC was strongly enriched and NFYA was also detected in the RBM25 interactome. Given the sequence-specific DNA-binding activity of NFYA (107), RBM25 may associate with the NFY complex rather than independently recognizing the CCAAT motif. Biochemical binding assays, such as electrophoretic mobility shift assays (EMSAs), together with domain- and motif-specific perturbations, will be needed to test direct DNA recognition by RBM25 and clarify its mechanism of chromatin recruitment. Beyond CEBPZ, our co-IP/MS analysis identified a diverse set of RBM25-associated proteins, reinforcing its multifunctional role in hESCs. While expectedly, many of these putative interactors are splicing factors, we also detected chromatin-associated factors such as CTCF and YY1 within the RBM25 interactome. Consistently, our CUT&RUN motif analysis revealed enrichment of CTCF-like motifs at intronic regions, suggesting a potential link between RBM25 and higher order chromatin organization, consistent with the previously reported role of RBM25 in YY1-mediated promoter–enhancer interactions (19). Another notable interactor observed in this study is SOX2, a core pluripotency factor known to repress neuronal lineage genes in hESCs (108). In our dataset, RBM25 binding is enriched in intronic regions of transcriptionally silent neuronal genes, which are predominantly located in closed chromatin. This suggests a potential cooperative role between RBM25 and SOX2 in maintaining repression of differentiation-associated genes. However, whether this interaction is direct and functionally significant remains to be determined. In addition, DNMT3B, an established de novo DNA methyltransferase involved in epigenetic regulation during development (109, 110), was identified among the top RBM25 interactors consistent with a recent study demonstrating that RBM25 KO reduces DNA methylation levels in mESCs (45). These observations indicate a potential link between RBM25 and DNA methylation–mediated gene regulation in human pluripotency, yet to be studied.

Collectively, our findings expand the functional landscape of RBM25 in hESCs. In addition to regulating RNA processing, RBM25 associates with chromatin and interacts with transcriptional and epigenetic regulators, suggesting potential roles in transcription, chromatin organization, and DNA methylation. These diverse interactions reinforce the role of RBM25 as a multifunctional regulator in hESCs, whose activity is shaped by its genomic context and interacting partners.

## Supplementary data

### List of supplementary figures

**Supplementary Figure S1-S8,** Supplementary Figures (PDF file).

### List of supplementary tables

**Supplementary Table S1.** ENCODE accession numbers and metadata for the histone-modification datasets used in this study. (Excel file)

**Supplementary Table S2.** List of ENCODE transcription factor and RNA-binding protein datasets selected for the TF–RBP chromatin co-occupancy analysis. (Excel file)

**Supplementary Table S3.** Results of TF-RBP co-occupancy analysis. (Excel file)

**Supplementary Table S4.** RBM25 eCLIP-seq peak annotations. (Excel file)

**Supplementary Table S5.** RBM25 CUT&RUN peak annotations. (Excel file) **Supplementary Table S6.** RBM25 nuclei interactome. (Excel file)

**Supplementary Table S7.** CEBPZ CUT&RUN peak annotations. (Excel file)

**Supplementary Table S8.** Differentially expressed genes and associated statistics following CEBPZ or RBM25 knockdown. (Excel file)

## Acknowledgements

We would like to thank Amir Argoetti for advice on the eCLIP experiment, Oded Kleifeld for his advice and help with proteomics analyses. We also thank Hiba Hassanain and Ira Rakita for sharing analysis code and providing helpful advice. We thank the Azrieli Technion Genomics Center for sequencing services and the Smoler Proteomics Center for LC–MS/MS data acquisition.

## Author Contributions

Y.S: Investigation, Formal analysis, Validation, Visualization, Writing – original draft. T.L: Formal analysis, Data curation. Writing – review & editing. F.G: Formal analysis, Visualization. T.H: Investigation, Validation, Writing – review & editing. Y. M-G: Conceptualization, Funding Acquisition, Supervision, Validation, Writing – original draft, Writing – review & editing.

## Data availability

The eCLIP-seq and CUT&RUN sequencing data have been deposited in the Gene Expression Omnibus (GEO) under accession numbers GSE343821 and GSE344038, respectively. All RNA-seq datasets used in this study are available in GEO under accession number GSE343820. The RBM25 interactome mass spectrometry data have been deposited with the ProteomeXchange Consortium under project accession PXD083004.

