## Supplementary Figures S1-S8 for "RBM25 exhibits distinct RNA- and chromatin-associated target programs in human embryonic stem cells"

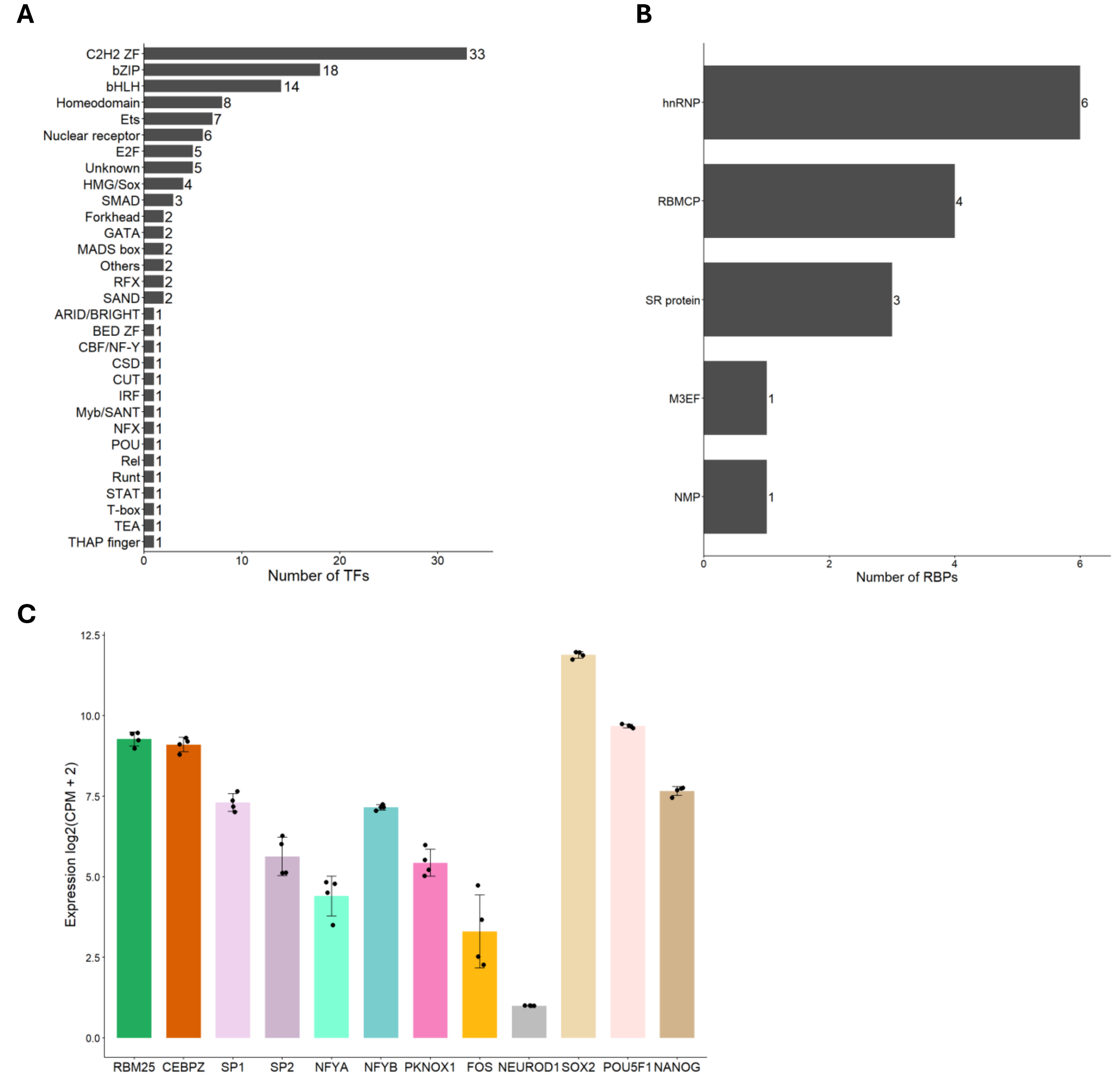


**Figure S1 Selected TFs and RBPs.**
**A.** Selected TFs used in co-occupancy analysis, classified by DBDs.
**B.** Selected RBPs used in co-occupancy analysis, classified by RBP families. SC: spliceosome component; M3EF: mRNA 3’end formation; RBMCP: RNA-binding motif containing protein; NMP: nuclear matrix protein.
**C.** Expression levels of RBM25 and high-Z-score RBM25-associated TFs in hESCs, with SOX2, POU5F1, and NANOG included as pluripotency-marker references.

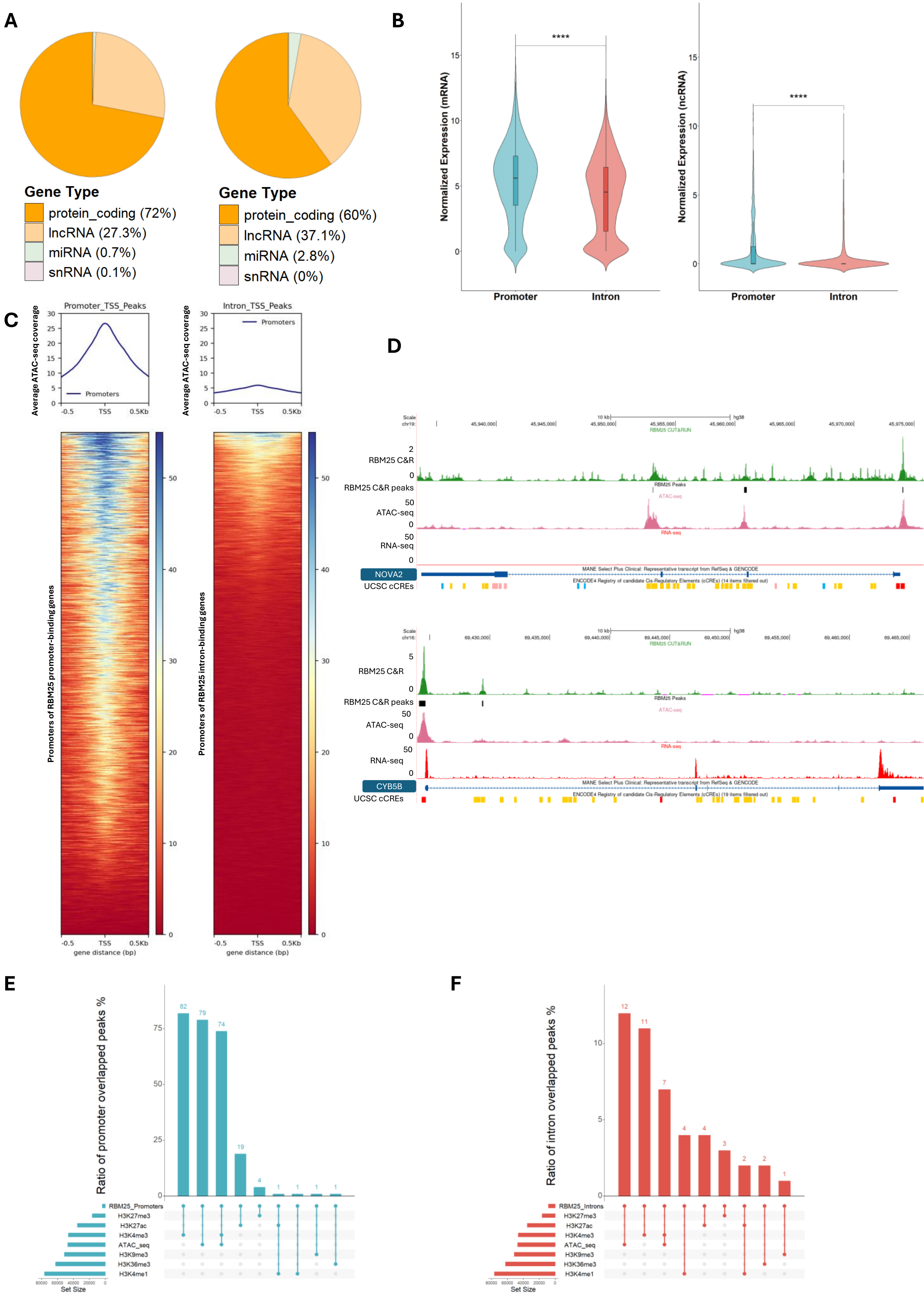


**Figure S2: Chromatin features associated with promoter- and intron-bound RBM25 target genes.**

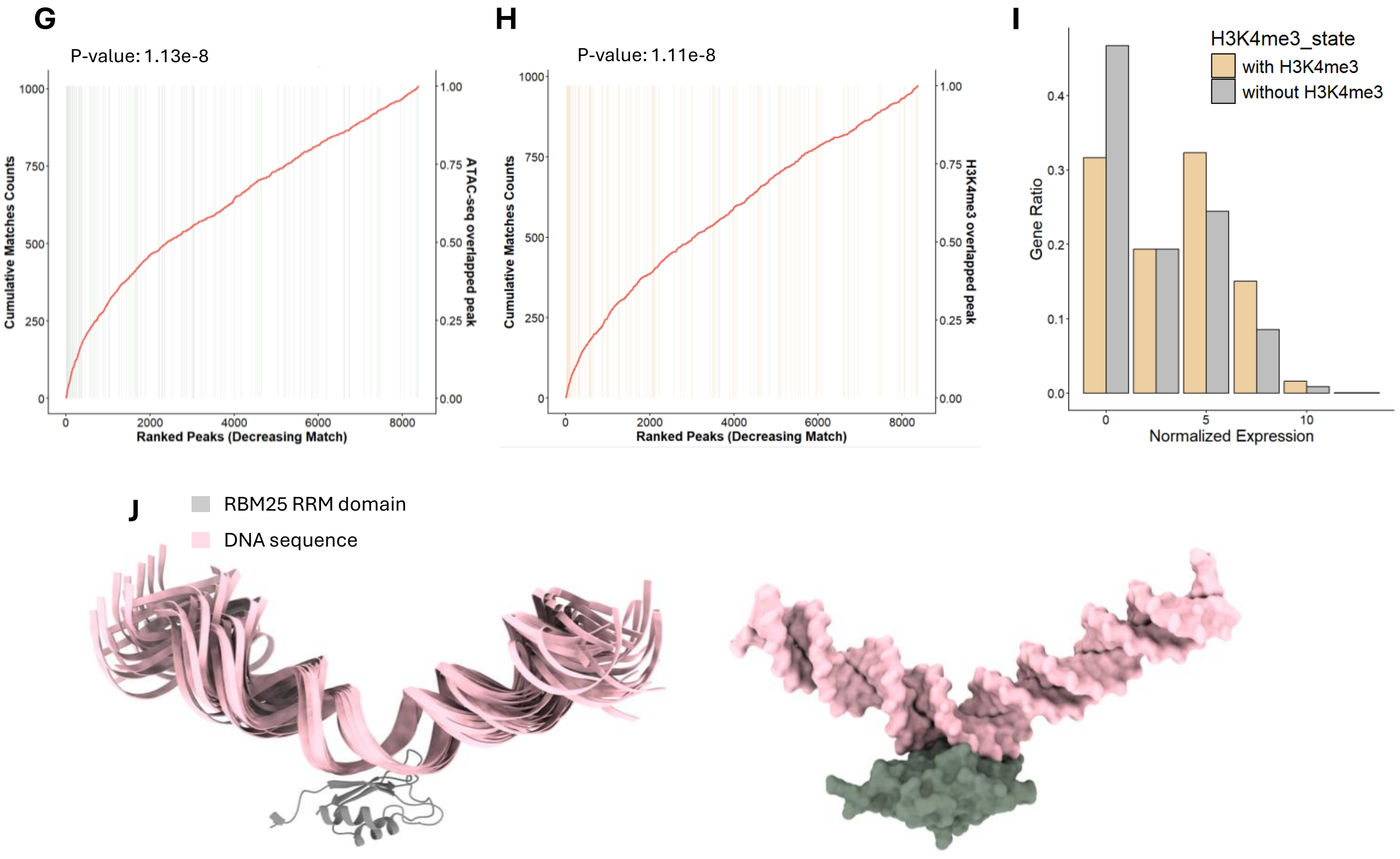


**Figure S2 (continued): Chromatin features associated with promoter- and intron-bound RBM25 target genes.**
**A.** Gene biotype distribution of RBM25-bound genes. Left, genes with promoter-localized RBM25 peaks. Right, genes with intronic RBM25 peaks.
**B.** Violin plots comparing normalized expression levels of promoter-bound and intron-bound genes after stratification by gene biotype. Left, protein-coding genes. Right, non-coding genes. Statistical significance was assessed by t-test.
**C.** ATAC-seq signal at promoters of RBM25 target genes. Left, genes with promoter-only RBM25 binding. Right, genes with intron-only RBM25 binding. Promoters were defined as ±500 bp from the transcription start site.
**D.** UCSC Genome Browser views showing RBM25 CUT&RUN signal, RBM25 peaks, ATAC-seq signal, and RNA-seq coverage at genes containing both promoter and intronic RBM25 peaks. Top, NOVA2. Bottom, CYB5B.
**E.** UpSet plot showing the overlap between promoter-localized RBM25 peaks and ATAC-seq or histone modification peaks.
**F.** UpSet plot showing the overlap between intronic RBM25 peaks and ATAC-seq or histone modification peaks.
**G.** Cumulative enrichment analysis of ATAC-seq across ranked intronic RBM25 peaks. Peaks were ranked by CUT&RUN peak score and analyzed using minimum hypergeometric enrichment analysis. mHG p-value is shown in the top left corner.
**H.** Cumulative enrichment analysis of H3K4me3 across ranked intronic RBM25 peaks. Peaks were ranked by CUT&RUN peak score and analyzed using minimum hypergeometric enrichment analysis. mHG p-value is shown in the top left corner.
**I.** Expression status of RBM25 intron-bound genes stratified by overlap between intronic RBM25 peaks and H3K4me3 peaks.
**J.** AlphaFold 3 model of the protein-DNA complex between the full RBM25 protein and a selected DNA sequence (see methods), shown is the RRM domain only that was predicted to interact with the DNA in the top scored models. Left, ribbon representation showing the superposition of 10 highly scored models. The RBM25 RRM domain is shown in grey and DNA in pink. Right, a surface representation of a selected model of the RBP25 RRM in complex with DNA. Predicted structure of the complex was visualized using UCSF Chimera (see methods)

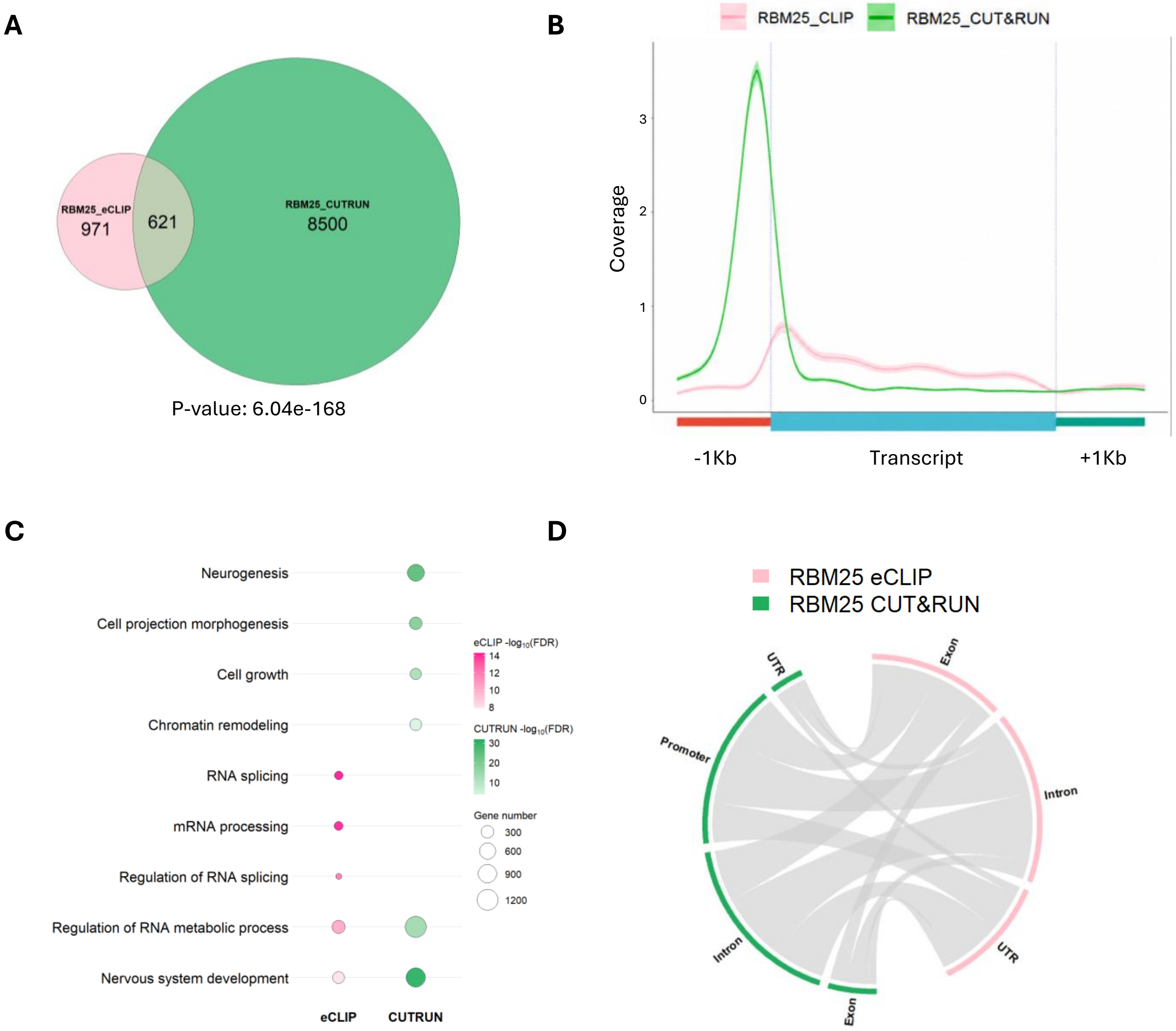


**Figure S3: Comparison of RBM25 RNA- and chromatin-associated target genes identified by eCLIP and CUT&RUN.**
**A.** Overlap between RBM25 target genes identified by eCLIP and CUT&RUN.
**B.** Metagene coverage profiles comparing RBM25 RNA-binding sites identified by eCLIP and RBM25 chromatin-binding sites identified by CUT&RUN across gene bodies. Pink, eCLIP signal. Green, CUT&RUN signal.
**C.** Gene Ontology enrichment analysis (for biological processes) of RBM25 target genes detected specifically by eCLIP, CUT&RUN, or both assays.
**D.** Positional comparison of RBM25 RNA-binding and chromatin-binding sites within genes bound both at the RNA and chromatin levels.

**A**

| RBM25 DNA motif | p-value | Similar Known motif |
|---|---|---|
| 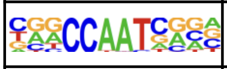 | $1 \times 10^{-524}$ | NFYA, NFYB, NFYC, CEBPZ |
| 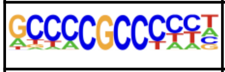 | $1 \times 10^{-274}$ | SP1, SP2 |
| 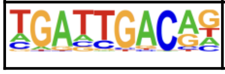 | $1 \times 10^{-62}$ | PBX3 |

| RBM25 DNA motif | p-value | Similar Known motif |
|---|---|---|
| 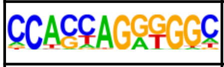 | $1 \times 10^{-103}$ | CTCF-like |
| 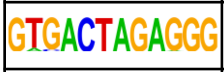 | $1 \times 10^{-16}$ | SMAD3 |
| 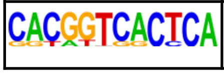 | $1 \times 10^{-10}$ | PKNOX1 |

**B**

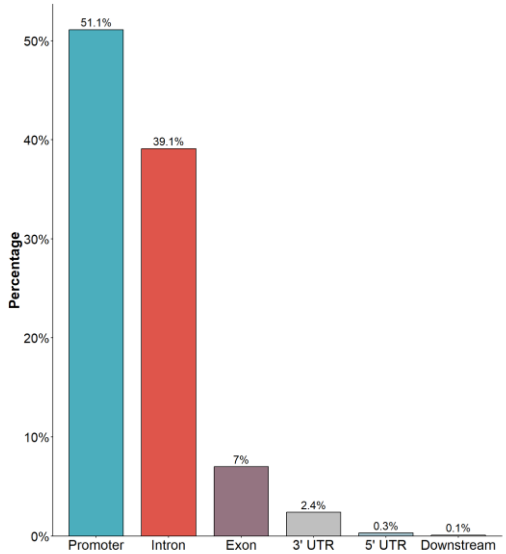


**C**

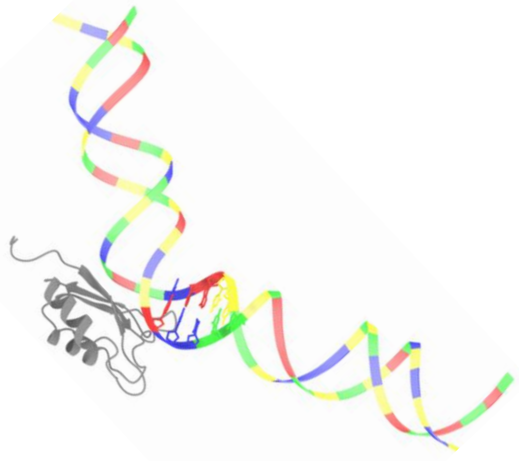

**D**

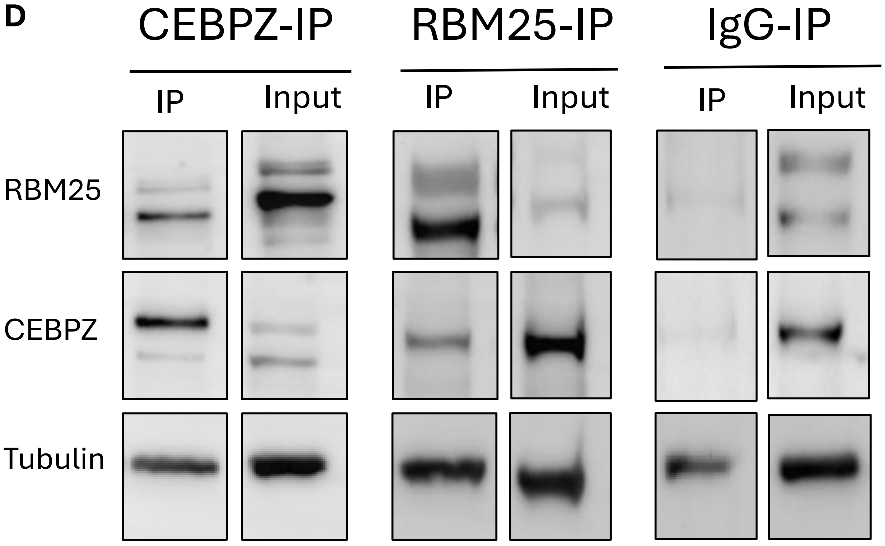


**Figure S4: Additional evidence supporting the association between RBM25 and CEBPZ.**
**A.** De novo sequence motifs enriched within promoter- and intron-associated RBM25 CUT&RUN peaks, identified using HOMER, together with the top-matched known transcription factor motifs. Left, promoter-associated RBM25 peaks. Right, intron-associated RBM25 peaks.
**B.** Genomic distribution of CCAAT-containing RBM25 CUT&RUN peaks.
**C.** AlphaFold 3-predicted structure of the RBM25 protein in complex with a double strand DNA sequence extracted from the CUT&RUN results, containing a CCAAT motif in the center (DNA sequence is provided in the method section). Displayed is the RRM domain (in grey), and the DNA predicted structure, colored according to nucleotide (A, red; T, blue; C, yellow; G, green). The CCAAT nucleotides positioned in proximity to RRM loops are highlighted. Predicted structure of the complex was visualized using UCSF Chimera (see methods)
**D.** Western blot validation of reciprocal co-immunoprecipitation between CEBPZ and RBM25 using whole-cell lysates. Tubulin is shown as a control.

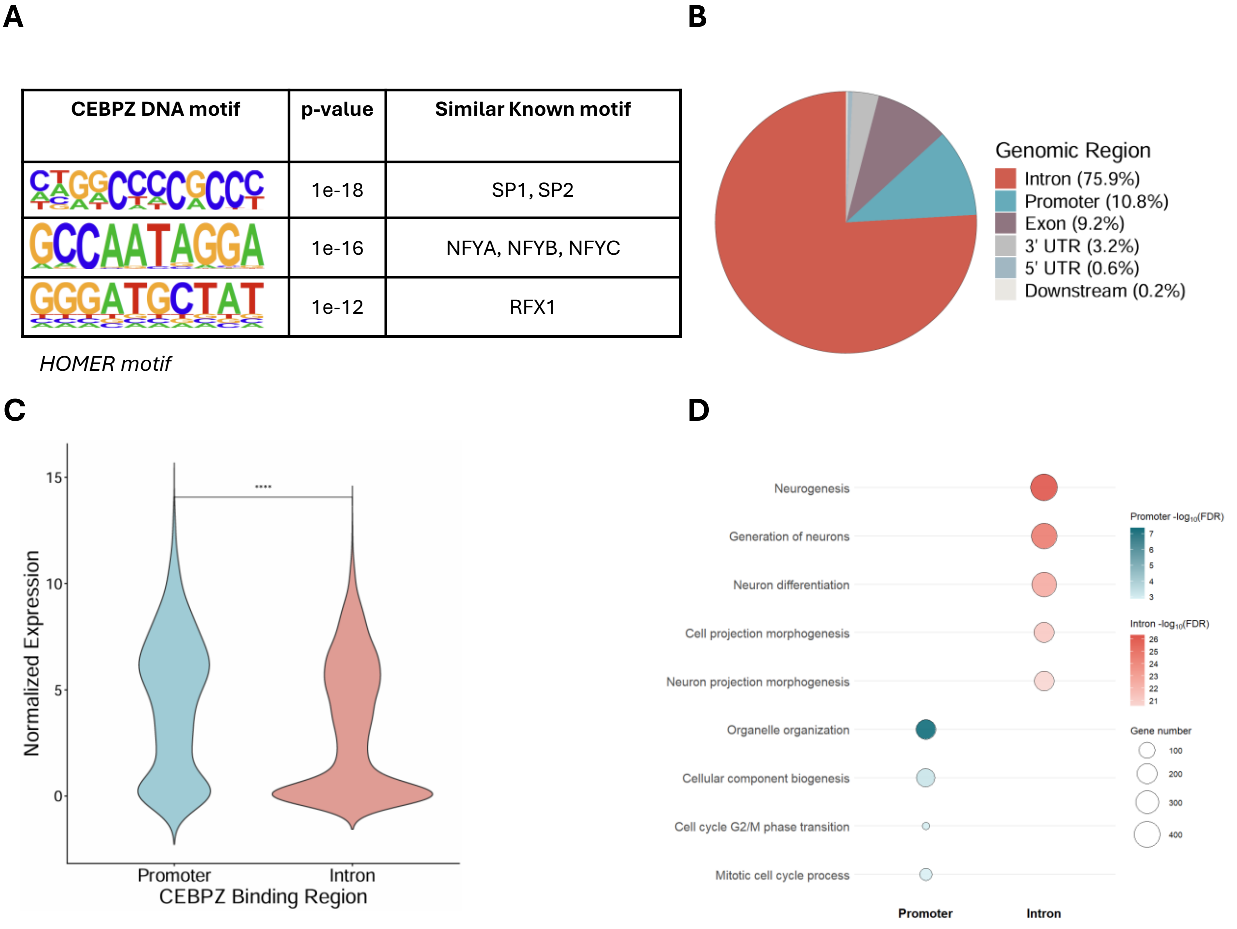


**Figure S5: Chromatin-binding features of CEBPZ in hESCs.**

**A.** De novo motifs enriched within CEBPZ CUT&RUN peaks, identified using HOMER, together with the top-matched known transcription factor motifs.

**B.** Genomic distribution of CEBPZ CUT&RUN peaks based on absolute peak counts.

**C.** Violin plot comparing normalized expression levels of genes with promoter-associated or intronic CEBPZ binding. Statistical significance was assessed using t-test.

**D.** Gene Ontology enrichment analysis (for biological processes) of genes with promoter-associated or intronic CEBPZ binding.

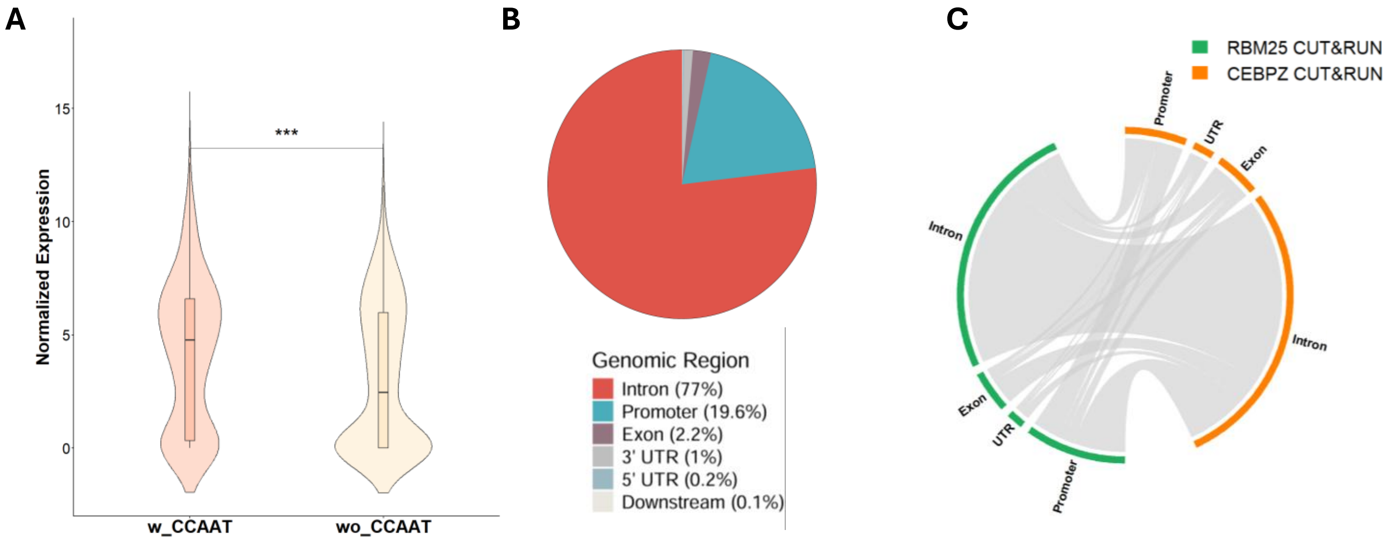


**Figure S6: Additional features of RBM25 and CEBPZ chromatin co-occupancy.**
**A.** Comparison of normalized expression levels between genes bound by bothRBM25 and CEBPZ, with or without a CCAAT motif within the corresponding peak.
**B.** Genomic distribution of regions bound by both RBM25 and CEBPZ.
**C.** Chord diagram showing the relationships between RBM25 and CEBPZ binding-region classes among genes bound by both proteins.

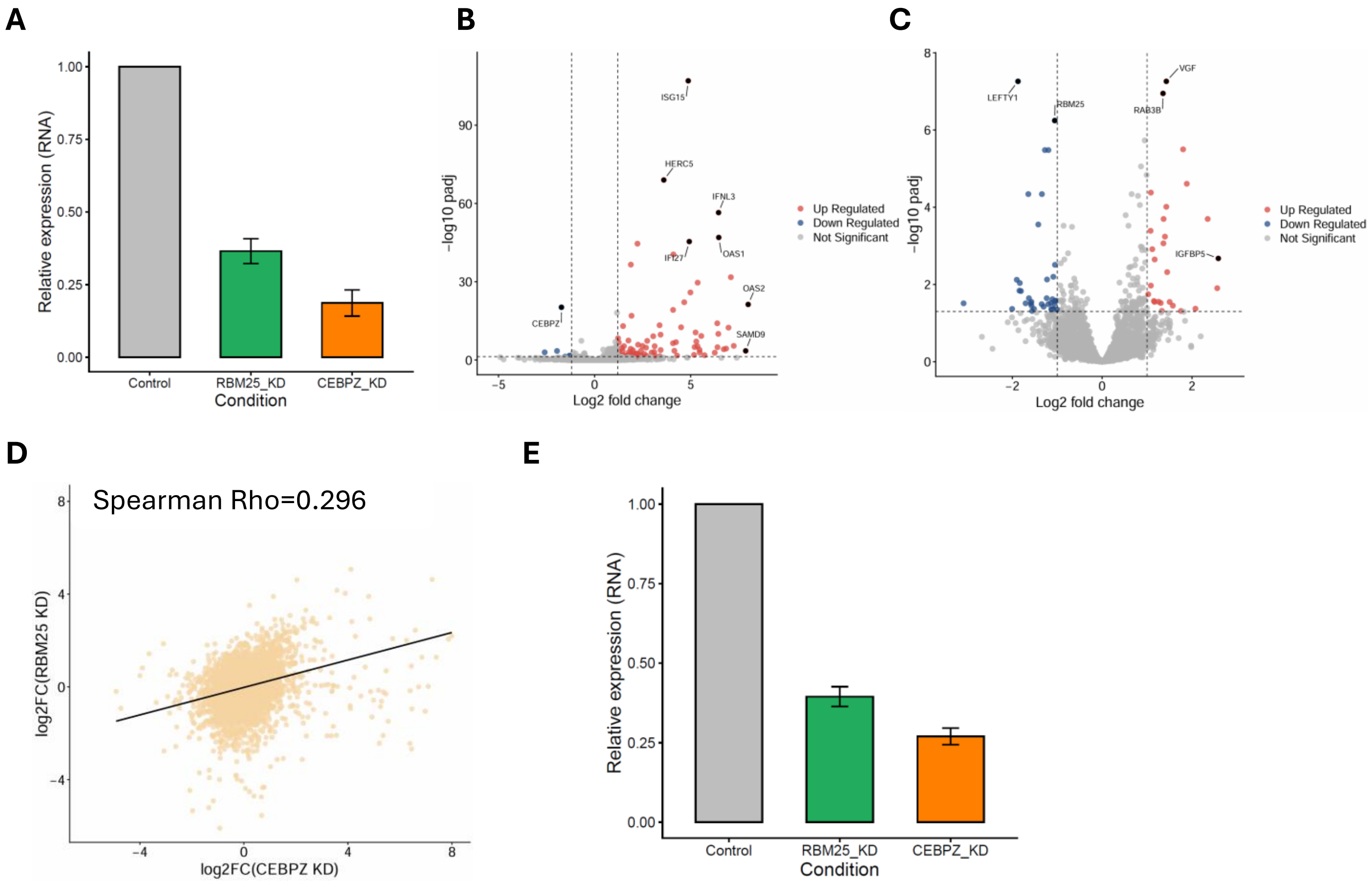


**Figure S7: Transcriptional effects of individual and combined RBM25 and CEBPZ depletion.**
**A.** Relative RBM25 and CEBPZ mRNA levels following individual siRNA-mediated KD, measured by quantitative PCR.
**B.** Volcano plot showing differential gene expression following CEBPZ KD.
**C.** Volcano plot showing differential gene expression following RBM25 KD.
**D.** Correlation between gene expression changes following RBM25 and CEBPZ KD. Log2 fold changes were compared using Spearman's correlation analysis.
**E.** Relative RBM25 and CEBPZ mRNA levels following combined KD, measured by quantitative PCR.

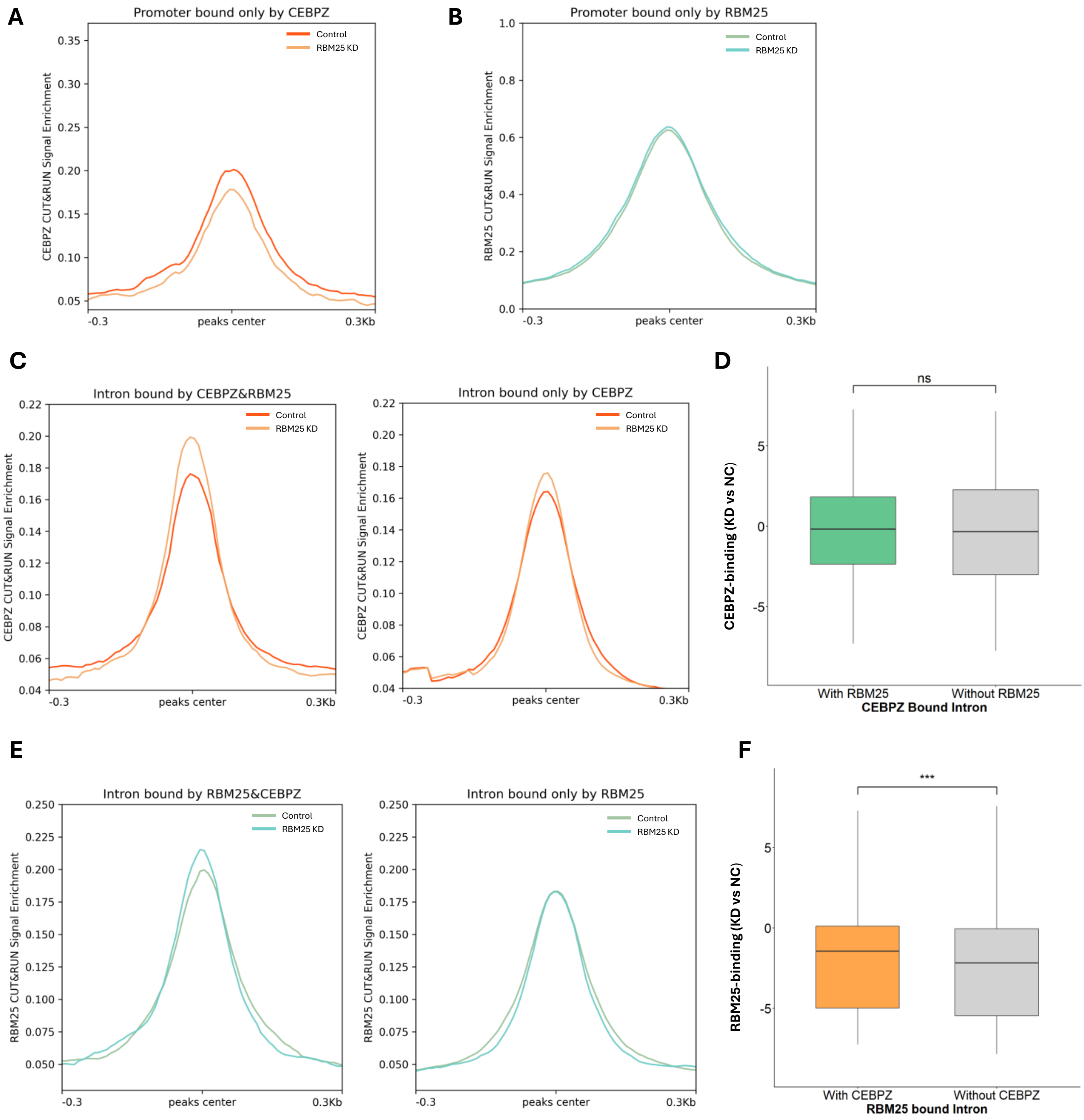


**Figure S8: Effects of reciprocal depletion on RBM25 and CEBPZ chromatin occupancy.**

**A.** Metagene profiles of CEBPZ CUT&RUN signal across promoter regions bound only by CEBPZ in control and RBM25-depleted cells.

**B.** Metagene profiles of RBM25 CUT&RUN signal across promoter regions bound only by RBM25 in control and CEBPZ-depleted cells.

**C.** Metagene profiles of CEBPZ CUT&RUN signal across intronic regions in control and RBM25-depleted cells. Left, introns bound by both RBM25 and CEBPZ. Right, introns are bound only by CEBPZ.

**D.** Distribution of log2 fold changes in CEBPZ CUT&RUN signal following RBM25 KD at genes bound by both RBM25 and CEBPZ and CEBPZ-specific intronic regions. Statistical significance was assessed using Student's t-test.

**E.** Metagene profiles of RBM25 CUT&RUN signal across intronic regions in control and CEBPZ-depleted cells. Left, intronic regions co-bound by RBM25 and CEBPZ. Right, intronic regions bound only by RBM25.

**F.** Distribution of log2 fold changes in RBM25 CUT&RUN signal following CEBPZ KD at co-bound and RBM25-specific intronic regions. Statistical significance was assessed using t-test.
